# DiscERN: An Automated Genome Mining Tool for the Discovery of Evolutionarily Related Natural Products

**DOI:** 10.1101/2025.09.10.675446

**Authors:** Jeremy G. Owen, Ethan F. Woolly, Hung-En Lai, V. Helen Woolner, Rory F. Little

## Abstract

Targeted genome mining to expand known families of natural products is a powerful strategy for discovering bioactive compounds, yet it remains a significant bioinformatics challenge. While tools exist for de novo biosynthetic gene cluster identification and large-scale unsupervised clustering, dedicated methods for the targeted, hypothesis-driven expansion of user-defined BGC families are lacking. Here, we present DiscERN (Discoverer of Evolutionarily Related Natural products), a userfriendly tool designed to address this gap. DiscERN leverages a multi-modal ensemble method that integrates four complementary algorithms classifying biosynthetic gene clusters based on Pfam content, sequence homology, and predicted product structure. This approach allows users to strategically balance discovery sensitivity with predictive precision to suit diverse research goals. We demonstrate DiscERN’s utility by applying it to a large collection of actinomycete genomes and validating its predictive power through the successful isolation of discomycin A, a new calcium-dependent lipopeptide antibiotic, from a silent biosynthetic gene cluster. DiscERN provides a robust and accessible platform that streamlines the path from genomic data to a prioritised list of candidate biosynthetic gene clusters, effectively bridging the gap between in silico prediction and bioactive compound discovery.

## Introduction

The inexorable rise of antimicrobial resistance constitutes a global health crisis, threatening to undermine pillars of modern medicine.^1^ In response, there has been a renewed search for new chemical scaffolds with antibacterial activity. Historically, bacterial natural products have been the most prolific source of antibiotics.^2^ These complex compounds are the result of millions of years of evolution, fine-tuning them for potent and specific biological activities.^3-5^ It is now widely recognized that the true biosynthetic potential of the microbial world remains vastly underexplored, offering a rich, untapped reservoir of chemical diversity for drug discovery and biomedical research.^6, 7^

Genomic data has revealed that the capacity to produce natural products is encoded within discrete, co-located sets of genes known as Biosynthetic Gene Clusters (BGCs),^8, 9^ where each BGC acts as a molecular assembly line, containing the complete enzymatic “blueprint” for constructing a specific natural product.^10^ The advent of high-throughput genome sequencing, coupled with advances in synthetic biology and heterologous expression, have revolutionised our ability to translate these genomic blueprints into chemical reality.^11-13^ Strikingly, genome mining studies consistently show that the number of BGCs identified in microbial genomes far exceeds the number of known compounds isolated from those same organisms.^6, 8^ This vast collection of “silent” or cryptic BGCs represents an opportunity for discovering new molecules, reducing the reliance on bioactivity guided screening, which is stochastic and frequently confounded by the rediscovery of known compounds.^2^

Computational tools are central to unlocking this opportunity. The development of antiSMASH,^14^ a widely-used platform for BGC annotation, and its rich informatics ecosystem have enabled researchers to systematically identify and classify BGCs from genomic data.^11, 15-17^ One of the most fruitful strategies in genome mining is the targeted expansion of known natural product families.^18-22^ BGCs responsible for producing a particular class of compounds, such as the glycopeptide antibiotics^23^ or the 16-membered polyketide macrolides,^24^ often share a conserved biosynthetic functions and sequence homology. By using known, well-characterised BGCs as a query, researchers can mine genomic datasets for evolutionary relatives that might produce new compounds. This genomics-driven approach has proven remarkably successful, leading to the discovery of novel structural variants with improved properties,^25^ distinct biological targets,^18, 20, 22^ or antimicrobials with activity against strains resistant to existing clinically employed relatives.^21, 22, 26^

While this targeted strategy is powerful, it has been hampered by a lack of dedicated, user-friendly tools. Existing state-of-the-art platforms like BiG-SCAPE^16^ and BiG-SLiCE^15^ are designed for the valuable but distinct task of *unsupervised* clustering. They excel at organizing massive BGC collections into similarity networks, providing a global overview of biosynthetic diversity. However, they are not optimized for a hypothesis-driven workflow where a researcher wishes to ask a specific question: “Given this defined set of BGCs for my family of interest, where can I find more like them?” Consequently, researchers are often forced to develop ad hoc search strategies for each family of interest, a process that typically involves manually scripting disparate tools and lacks efficiency, scalability, and reproducibility.

To address this challenge, we developed DiscERN (Discoverer of Evolutionarily Related Natural products), an automated and robust method for the targeted expansion of user-defined BGC families (**Figure 1**). Recognising that evolutionary relatedness can be captured through different but complementary features,^3-5, 27^ we built DiscERN around an ensemble of four distinct algorithms. These approaches classify BGCs based on three principles: (i) overall Pfam domain content, providing an architectural fingerprint; (ii) direct protein sequence similarity; and (iii) predicted structure of the final small-molecule product, offering a functional chemical perspective. By integrating these methods into a flexible and intuitive pipeline, DiscERN provides an efficient workflow for analysing large BGC datasets to expand the diversity of target compound classes. As a demonstration of its utility, we applied our method to a collection of 3561 actinomycete genomes. This search identified numerous BGCs putatively encoding new members of existing natural product families. We activated one of these using heterologous expression and regulatory gene overexpression, leading to the isolation discomycin A, a new member of the calcium-dependent lipopeptide family from a silent BGC in the genome of *Streptomyces kanamyceticus* ATCC12853.

**Figure 1.**
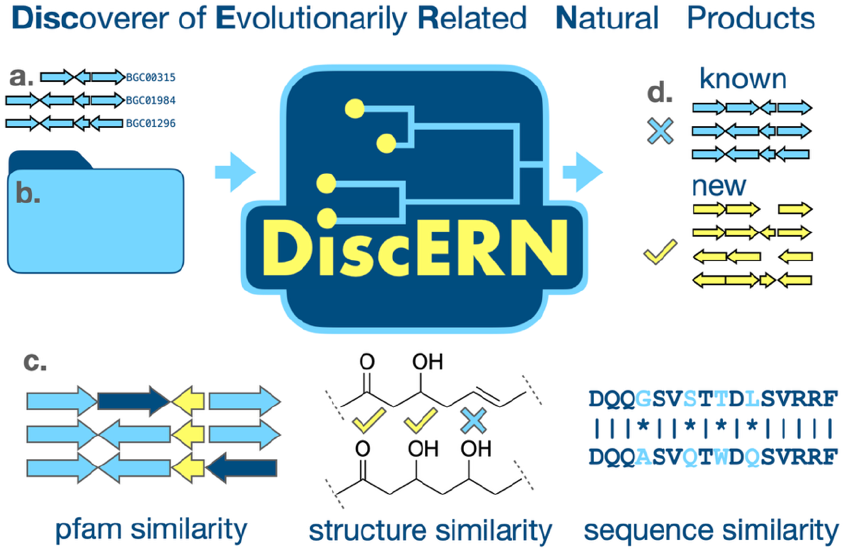
Schematic overview of DiscERN workflow: A user defined family of BGCs specified using MiBiG numbers (**a**) and directory containing antiSMASH outputs for genomes to be analysed (**b**) are provided as inputs. DiscERN then applies and integrates an ensemble of algorithmic approaches to identify new family members (**c**) and provides Intuitive outputs (**d**) to facilitate prioritisation for downstream functional analysis.

## Results

### Development of Complementary Algorithms for BGC Family Expansion

To create a robust tool for targeted BGC family expansion, we developed four complementary classification methods designed to capture different aspects of evolutionary relatedness. The first of these is the **Pfam Vector (pfam-vec)** method, which adapts the BGC vectorization approach from BiG-SLiCE,^15^ and classifies BGCs based on their overall Pfam domain content. This method converts antiSMASH outputs into numerical vectors representing the presence, abundance, and amino acid sequences of Pfam domains, providing a high-level architectural fingerprint of each BGC. To provide a reference for development and benchmarking, we defined six manually curated antibiotic families from the MiBiG database^17^ (**Supplementary Data Table S1**): calcium-dependent lipopeptides^28^, glycopeptides,^23^ macrolides,^24^ aminoglycosides,^29^ rifamycin like polyketides^26, 30, 31^ and polyene antifungals.^22, 32^ For each family, we calculated a centroid vector representing the average Pfam domain composition and determined an optimal cosine distance threshold to classify family members against a background of all other BGCs, with model performance assessed using the Matthews Correlation Coefficient (MCC).^33^ The maximum achievable MCC score varied (0.8 to 1.0), suggesting some families are more easily distinguished by Pfam content than others (**Figure 2**).

**Figure 2.**
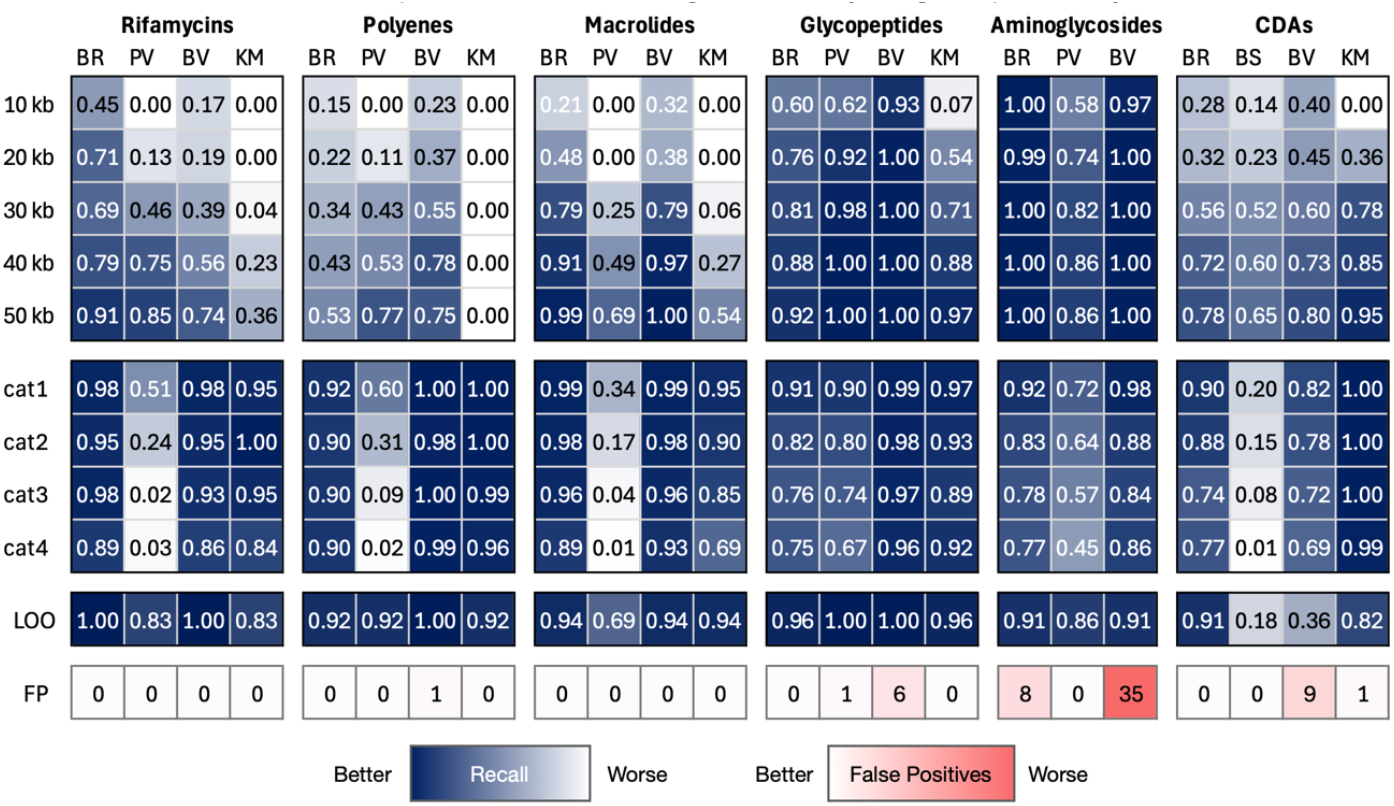
Evaluation of performance for each classification method: Heat maps indicate performance of each method in classification of BGCs belonging to six manually defined antibiotic families. Each panel represents a single antibiotic family. Column labels indicate classification methods: BR = blast-rank, BV = blast-vec, PV = pfam-vec, KM = structural k-mer intersection. Row labels indicate assessment metrics: 10- 40 kb = Sensitivity of approach when applied to 100 randomly generated BGC sub-fragments of the indicated size. Cat1-4 = Sensitivity of approach when applied to 100 BGCs concatenated with 1-4 additional randomly selected BGCs. LOO = Sensitivity of approach in leave-one-out benchmarking. FP=total false positives among 2613 BGCs extracted from 150 randomly selected actinomycetes genomes.

The centroids and optimal cosine distances determined for each BGC family comprise a model that can be used for classification. In our workflow, pfam-vecs are generated for all BGCs to be classified, and their cosine distance from a target BGC family centroid calculated, with BGCs below the previously determined optimal threshold classified as belonging to the family in question. A leave-one-out analysis using this approach to assess out-of-train sensitivity showed variable performance among BGC families. While glycopeptides performed perfectly (recall=1.0) and others reasonably well (rifamycins, polyenes, aminoglycosides; recall=0.83-0.92), the macrolide model was average (recall=0.69) and the CDA model exhibited poor performance (recall=0.18) (**Figure 2, Supplementary Data Table S2**). When challenged with a test set of five recently discovered CDAs not present in the training data,^20, 34-37^ the model’s limited generalisability was confirmed, as it correctly identified only three of the five test cases.

### Evaluation of robustness to sequencing artefacts

We further evaluated robustness by simulating fragmented BGCs and concatemers of adjacent BGC, common artifacts arising from fragmented genome assemblies and overlapping BGC boundaries, respectively.^38, 39^ We generated concatemers by appending a randomly selected BGC (from the antiSMASH 7.0 output of *Actinomycetes* genomes) to each of the MiBiG reference BGCs in our chosen antibiotic families. Fragments were generated by extracting random sequence segments of defined lengths (10, 20, 30, 40, and 50 kb for glycopeptides, lipopeptides, and macrolides; 10 and 20 kb for the smaller aminoglycoside clusters) from the reference BGCs and re-analysing them with antiSMASH.

For each reference BGC, we generated 100 concatemers, as well as 100 fragments of each specified size. These simulated fragmented and concatenated BGCs were then analysed using the models trained for each compound class. The models performed reasonably well on larger fragments but exhibited significantly reduced performance on smaller fragments and concatemers (**Figure 2**). This highlighted a need for an alternative scoring method to improve sensitivity in these challenging scenarios.

### Evaluation of Specificity

To evaluate the specificity, we compiled a BGC collection by analysing 150 randomly selected *Actinomycetes* genomes using antiSMASH version 7.0,^14^ resulting in the annotation of 2613 BGCs. We then applied the models generated using each of our six manually curated antibiotic BGC families to this collection. To our knowledge, there are no existing automated tools that perform the task of targeted expansion of user defined BGC families, so manual analysis conducted by experienced practitioners was the best method for defining ground-truth and subsequently calculating false positive rate. A total of seven hits were returned across all six BGC families. Only one of these was deemed to be a false positive, indicating relatively high specificity for all six families tested.

### Development and Performance of Sequence Similarity Classifiers

To improve sensitivity, especially for fragmented BGCs and out-of-sample cases, we developed two further classifiers based on direct sequence similarity. These were assessed for sensitivity, specificity and robustness to sequencing artefacts as previously described (**Figure 2**).

The **“blast-vec”** method creates “blast-vectors” by treating the cumulative BLAST score of a query BGC against each MiBiG reference as a dimension. Using the same centroid-based approach, this method improved sensitivity but led to a reduction in specificity compared to the Pfam-based method. (**Figure 2**).

The **“blast-rank”** method uses a rank-based similarity metric. It defines a family based on the number of non-self, in-family BLAST hits (*n*) found within a certain rank (*k*) for all known family members. A query BGC is then classified as a hit if it has an in-family BGC as its top hit and at least *n* in-family hits within the top *k* results. The blast-rank method demonstrated improved out-of-train sensitivity and greater robustness to fragmented and concatenated BGCs, with only a slight reduction in specificity compared to the Pfam vector method. Its performance on fragmented data was comparable to or better than the blast-vector method (**Figure 2**).

### Development and Performance of the Predicted Product Structure Classifier

For NRPS/PKS families, we also developed the **“structural k-mer intersection”** method. This method parses antiSMASH JSON outputs to extract a linearized monomer sequence for each ORF in a BGC. These sequences are converted into unique k-mer feature sets. For classification, a query BGC’s k-mer set is compared against reference families, and it is assigned to the family with the best k-mer overlap, provided the score exceeds a pre-optimized threshold. This approach performed well, with recall values ranging between 0.82 and 0.96 for the five antibiotic families tested in our leave-one-out analysis (**Figure 2**). Specificity was also high, with just one false positive returned across all five antibiotic families when tested using our test collection of 150 actinomycetes genomes (2613 BGCs). The k-mer intersection method performed relatively well when tested using concatemers of target BGCs. In contrast, the method’s performance degraded significantly on smaller BGC fragments (**Figure 2**). This was expected, as fragmented clusters often lack the complete set of megasynth(et)ase modules required to reconstruct a meaningful k-mer profile, thereby precluding accurate classification based on product structure.

### Development of a Flexible Ensemble Method for Intuitive Automated Analysis

Evaluation of the algorithms we developed for BGC classification was confounded by the lack of a complete ‘gold standard’ dataset with known true negatives and false negatives, a common issue in the development of novel bioinformatics tools. The MiBiG database provides an excellent source of characterised BGCs to serve as labelled training data, however the relative scarcity of characterised BGCs means that partitioning this into test/train sets for cross validation is not feasible. Therefore, we evaluated the effectiveness of the component algorithms that were subsequently combined in DiscERN using a multi-faceted approach focused on the metrics most relevant to a real-world discovery scenario: sensitivity to known family members using leave-one-out, robustness to common data artifacts, and the specificity of predictions on an uncharacterized dataset. The four approaches we developed had variable performance across the six datasets we compiled and tested. In light of this, to leverage the strengths of the different approaches, we combined them into an ensemble method named DiscERN (<u>Disc</u>overer of <u>E</u>volutionarily <u>R</u>elated <u>N</u>atural products).

DiscERN is a command line application, implemented in python, that utilises a combination of all four classification methods along with an optional filtering step based on specific Pfam domain presence/absence. DiscERN allows users to define their own custom families of natural product BGCs for expansion by either specifying the MiBiG BGC numbers for each member, or by providing separate antiSMASH formatted GenBank and JSON files as a reference. It then scans user provided antiSMASH outputs for additional members of the BGC family and outputs a collection of combined GenBank files for the hits, grouped according to the number of algorithms (between 1-4) that independently identified a given hit. Hits are additionally segregated into those passing or failing a Pfam content filtering step that is automatically implemented based on common Pfam domains in the BGCs defining the family to be expanded. Outputs can then be optionally passed to a local install of antiSMASH for analysis. The primary outputs of DisCERN are a collection of genbank files, and optionally, collated antiSMASH outputs, that are grouped according to likelihood of truly representing on-target BGCs. These outputs are intended to facilitate selection of targets for heterologous expression or compound isolation studies. DiscERN can be installed for Linux and MacOSX and is available at https://github.com/MaxMeta/DiscERN.

### Performance of the DiscERN Ensemble on a Real-World Discovery Task

To assess performance in a realistic discovery scenario, we applied DiscERN to a large dataset of 3,561 publicly available Actinomycete genomes. To construct this dataset, we queried the NCBI Assembly Database^40^ for all genomes belonging to the class Actinomycetes and refined the resulting list to include only those designated as ‘representative’ or ‘reference’ genomes. Downloaded fasta files were then processed with antiSMASH 7.0 to yield 59,236 BGCs.

DiscERN was used to parse antiSMASH outputs for our collection of 3,561 genomes using four different antibiotic families as references: calcium-dependent lipopeptides, glycopeptides, rifamycin like polyketides and 16-membered macrolides. This search returned 688 putative hits, which were ranked based on the number of algorithms in the DiscERN ensemble providing support (a confidence score of k=1 to 4). To measure the precision of these confidence tiers, we manually curated hits for each family and tier to assign hits as either true-positives, false-positives. Hits not able to be determined owing to fragmentation were excluded from the analysis. This analysis revealed a clear trend: the precision of a prediction increased markedly with the level of ensemble agreement (**Figure 3, Additional Data File 2**). Predictions supported by at least three algorithms (k≥3) were highly precise, with a mean positive predictive value of 0.98. In contrast, precision was moderate for hits supported by two algorithms (k≥2, mean precision ∼0.6) and poor for those supported by only a single algorithm (k=1, mean precision <0.4), highlighting the power of the ensemble approach to stratify hits by confidence.

**Figure 3.**
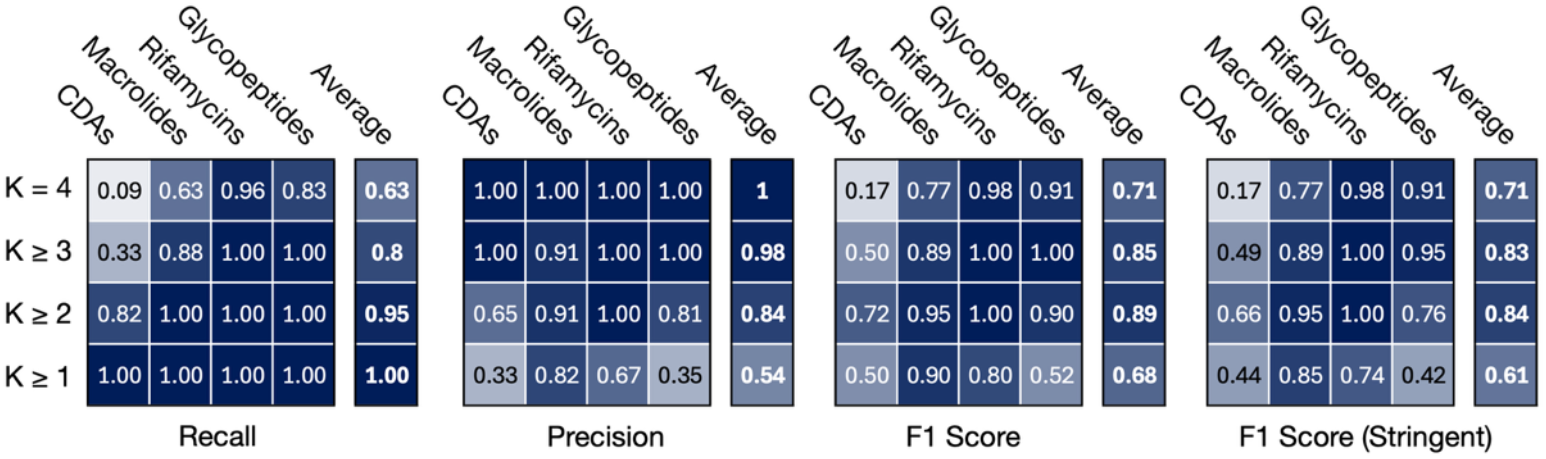
Performance of the DiscERN Ensemble: From left to right, heat maps indicate: Recall as assessed by leave one out cross validation; precision determined by manual examination of putative hits; F1-scores calculated from respective recall and precision values at each confidence tier; stringent F1 scores calculated by treating all indeterminant hits as false positives.

Owing to the large number of BGCs in our Actinomycete dataset, complete manual analysis to count false negatives was not feasible. In lieu of this, we estimated recall for four benchmark BGC families using the leave-one-out (LOO) cross-validation framework previously conducted for each of the algorithms individually (**Supplementary Data Table S2**). For each fold of the LOO analysis, the held-out BGC was classified by the full DiscERN ensemble, and the number of supporting algorithms (k) was recorded. We then calculated the cumulative recall for each confidence threshold (k≥1 to k=4) by determining the fraction of family members that were successfully identified at or above that level of ensemble support. This approach provides a robust estimate of the ensemble’s sensitivity and its ability to identify known true positives at different levels of stringency.

Our analysis allowed us to construct a comprehensive performance profile for DiscERN, balancing the precision observed in a real-world discovery scenario with the recall estimated from a curated dataset (**Figure 3, Supplementary Methods S3**). To summarise this trade-off, we calculated estimated F1 scores for each confidence level. The results show that a threshold of k≥2 achieves the highest F1 score (0.89), representing the optimal mathematical balance between a high recall (0.95) and strong precision (0.84). However, for many practical discovery applications where minimising experimental validation of false positives is a primary concern, a higher confidence threshold may be preferable. At k≥3, the precision is excellent (0.98), meaning nearly all predicted hits are true positives. This near-certainty is achieved while maintaining a robust recall of 0.80, resulting in a very strong F1 score of 0.85. This demonstrates that DiscERN can be tuned for different discovery goals: a threshold of k≥2 provides the optimal balance between precision and recall, while k≥3 offers a highly precise set of candidates that may be preferred if minimising downstream manual analysis is a priority. We also calculated a “worst case scenario” outcome, treating all hits classified as N/D as false positive. Even under this more stringent estimate DiscERN still performed well with average F1 scores of 0.83 and 0.84 at k≥3 and k≥2, respectively (**Figure 3**).

### Visualizing BGC Relatedness to Identify Novel Compound Candidates

DiscERN generates several outputs to facilitate hit prioritization. It calculates a mean pairwise distance matrix that integrates all three similarity measures (sequence similarity, Pfam content, and, where applicable, predicted product structure). This matrix, as well as the individual matrices for each distance metric, are used to construct dendrograms that intuitively display the degree of relatedness among the hit BGCs and the reference BGCs used to define the family. The combined matrix is also provided as a Newick formatted file for visualization in external tools like iTOL.^41^

When applied to the four BGC families from our Actinomycetes analysis, this feature allowed us to clearly identify distinct sub-clusters and outliers that were strong candidates for producing uncharacterised family members (**Figure 4, Supplementary Data Figure S14**). To further investigate these candidates, we compared the predicted NRPS/PKS domain compositions and substrate specificities of selected hits against their closest characterised relatives. We found 29 BGCs that clearly differed from their closest characterised relative, suggesting they encode novel compounds, and 85 BGCs that had identical domain composition and substrate specificity to their closest relative (**Figure 4, Additional Data File 1, Additional Data File 2**). This number is likely a conservative estimate of novelty, as this analysis does not account for variations in post-scaffold tailoring reactions, which are common in families like the rifamycins and glycopeptides.

**Figure 4.**
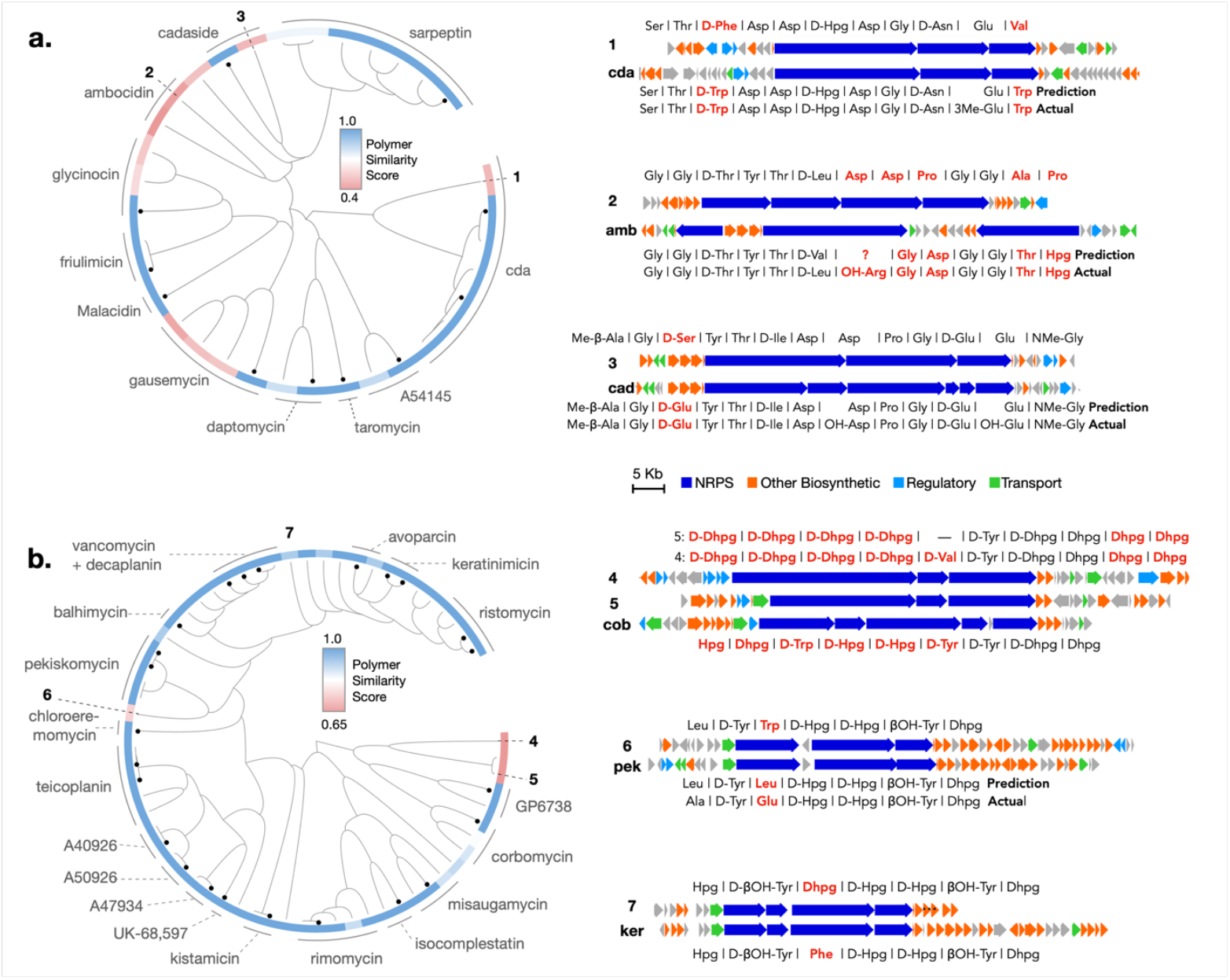
Visualisation of BGC relatedness using DiscERN: Putative hits for lipopeptide (**a**.) and glycopeptide (**b**.) antibiotics are shown. Left hand panels show hierarchical clustering results calculated by DiscERN using an average three distance metrics. The coloured bars at the end of each leaf indicate polymer similarity score, as determined by the kmer intersection algorithm implemented as part of the DiscERN ensemble. Black dots at leaves indicate the location of previously characterised reference BGCs that are used to define clades. Right hand panel shows selected BGCs predicted to encode new family members. Numbers match leaf labels. Predicted differences in NRPS specificity are highlighted in red.

### Discovery of the Discomycin BGC

Among the collection of BGCs identified as potentially encoding new bioactive compounds, a putative hit in the CDA family located within the genome of *S. kanamyceticus* ATCC12853 was chosen for further study (**Figure 4a**, BGC-1). This BGC, which we name the discomycin (*dsc)* BGC was chosen as it was present in a commercially available strain with no previous reports of calcium dependent antibiotic activity, and contained a SARP gene that suggested a facile means for activation.

The *dsc* BGC has clear similarity with the *cda* BGC from *S. coelicolor*. Both feature an 11-module NRPS architecture split across three contiguous genes. The *dsc* BGC contains the expected genes for for 4-hydroxyphenylglycine (Hpg) biosynthesis (*dscAJKL)* and glutamate 3-methylation (*dscV*). However, predicted A-domain specificities suggest the *dsc* BGC encodes a novel lipodepsipeptide, differing from CDA at residues 3, 9, and 11 (**Figure 4a, Figure 5a**).

**Figure 5.**
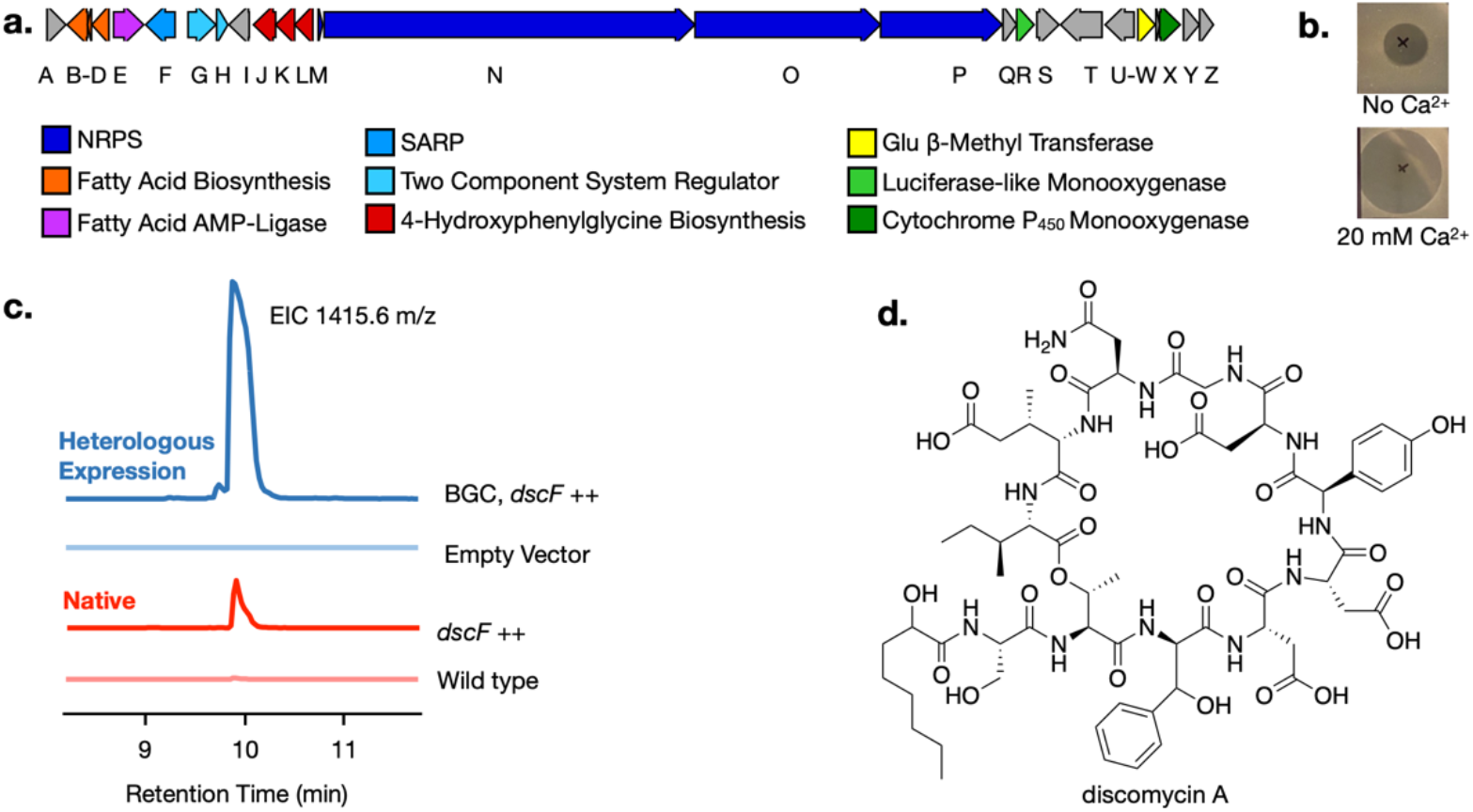
Discovery of Discomycin A using DiscERN: **A**. Map of the discomycin BGC predicted to encode a new calcium dependent lipopeptide antibiotic. **B**. Zone of inhibition assays showing calcium dependent antibiotic activity of crude extract from a culture of S. albus-del14 harbouring the discomycin BGC and an additional copy of the SARP gene *dscF* under the control of a strong constitutive promoter. **C**. LC-MS analysis of crude extracts from *S. kanamyceticus* and *S. albus*-del14 harbouring the discomycin BGC, each with an additional copy of the SARP gene *dscF* under the control of a strong constitutive promoter. **D**. Elucidated structure for discomycin A.

Like the *cda* BGC, the *dsc* BGC contains a dedicated Type II fatty acid synthase (FAS) system for synthesizing its lipid tail, including genes for chain initiation (*dscC*, KAS III), extension (*dscB*, KAS I), and an acyl carrier protein (*dscD*, ACP). The *dsc* cluster lacks the desaturase (*hxcO*) and epoxidase (*hcmO*) homologues required to produce the 2,3-epoxyhexanoyl moiety of CDA and possess a fatty acyl-AMP ligase (FAAL) gene, *dscE*. The presence of this FAAL suggests a hybrid lipidation mechanism, distinct from both the canonical CDA and daptomycin pathways. We hypothesize that the fatty acid is first synthesized by the on-pathway FAS system and then released, after which it is re-activated as an adenylate by the FAAL before being transferred to the NRPS assembly line.

### Activation of the Silent Discomycin BGC

To activate the silent *dsc* BGC, we first integrated an additional copy of the SARP gene, *dscF*, into the chromosome of the native producer, *S. kanamyceticus*, under the control of a strong constitutive promoter to give the strain *S. kanamyceticus::dscF*. Comparison of *S. kanamyceticus::dscF* to an empty vector control revealed the production of a new compound with a [M+H]^+^ ion at *m/z* 1415.6023, consistent with the expected mass of the predicted lipopeptide, which we named discomycin A (**Figure 4c**).

To definitively link this compound to the BGC, we cloned the entire *dsc* cluster using the CRISPR-Cas12a CAPTURE method^42^ and expressed it in the genome-minimized host *S. albus* Del14.^43^ This resulted in calcium dependent antibiotic activity in crude extracts from the resulting heterologus expression strain (**Figure 4b**) and LC-MS/MS analysis of this recombinant strain revealed an ion with an identical *m/z* and MS^2^ fragmentation pattern to that observed in the native host, confirming that discomycin A is the direct product of the *dsc* BGC.

Finally, to optimise production for isolation and structure elucidation, we introduced the *dscF* overexpression cassette into the heterologous host. The resulting strain, *S. albus* Del14::*dsc-dscF*, had higher production of discomycin A than either the heterologous host alone, *or S. kanamyceticus::dscF*, and was therefore selected for large-scale fermentation (**Figure 4b**)

### Isolation and Characterisation of Discomycin A

For structure elucidation, discomycin A was isolated from *S. albus* Del14::*dsc-dscF* grown in liquid TSB medium containing HP-20 resin. The compound was purified from the methanolic resin extracts of the harvested HP-20 resin using calcium-mediated precipitation followed by reversed-phase HPLC. Its planar structure was then elucidated using a combination of HRESI-MS/MS, 1D, and 2D NMR experiments (**Supplementary Data Table S5, Supplementary Data Figures S4-S13**).

The data revealed that discomycin A is a cyclic lipodepsipeptide containing 11 amino acids with an N-terminal 2-hydroxyoctanoyl tail. A 31-membered macrolactone ring is formed between the hydroxyl of Thr^2^ and the C-terminal carboxyl of Ile^11^. The elucidated peptide sequence—Ser^1^-Thr^2^-3OHPhe^3^-Asp^4^-Asp^5^-Hpg^6^-Asp^7^-Gly^8^-Asn^?^-3MeGlu^10^-Ile^11^—was in full agreement with the antiSMASH predictions for the *dsc* BGC. The structure of discomycin A was further supported by MS^2^ fragmentation of the NaOMe linearised the product (**Supplementary Data Figure S6)**

Finally, to determine the absolute stereochemistry of discomycin A, we performed C3 Marfey’s analysis^44^ on the hydrolysed peptide (**Supplementary Data Figure S7, Supplementary Data Table S6**). This confirmed the presence of D-Phe^3^, D-Hpg^6^, and D-Asn^?^, consistent with the epimerase domains predicted in the corresponding NRPS modules and allowed assignment of absolute configurations for all stereo centres except the β-carbon of 3-OH-Phe and C2 of the lipid tail.

### Spectrum of Activity of Discomycin A

The biological activity of purified discomycin A was evaluated using broth microdilution assays against a panel of Gram-positive and Gram-negative bacteria, alongside a cytotoxicity assessment against the HCT-116 human colon carcinoma cell line (**Supplementary Data Table S7**). Discomycin A exhibited potent, Ca^2+^-dependent antibacterial activity against a range of Gram-positive pathogens (**Figure 6a, Supplementary Data Table S7)**. with minimal inhibitory concentrations (MICs) in the low-micromolar range. In contrast, no antibacterial activity was observed against any of the Gram-negative bacteria in our panel, even at the highest tested concentration. Furthermore, discomycin A displayed no discernible cytotoxicity against HCT-116 cells, indicating a favourable selectivity for bacterial over mammalian cells. We also treated *S. aureus* with a sub-inhibitory concentration of compound. This lead to accumulation of N-acetylmuramic acid pentapeptide, indicating inhibition of cell wall biosynthesis (**Figure 6b**)

**Figure 6.**
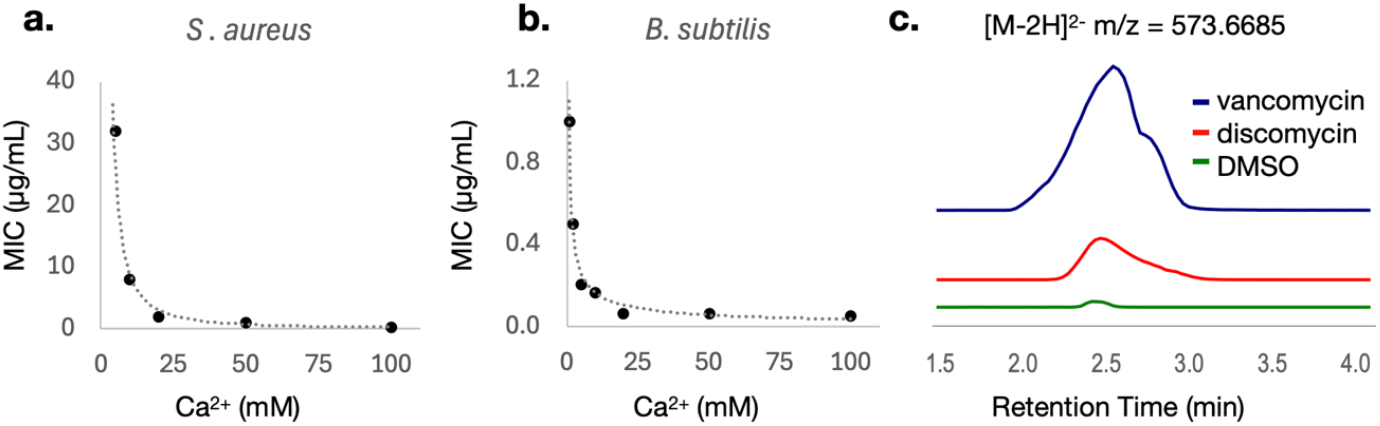
Discomycin has calcium-dependent antibiotic activity against Gram-positive bacteria and inhibits cell-wall biosynthesis: **A**. Average MIC values from triplicate assays against *S. aureus* at various concentrations of Ca^2+^. **B**. Average MIC values from triplicate assays against *B. subtilis* at various concentrations of Ca^2+^. **C**. HPLC-HRMS analysis of cell lysate from *S. aureus* cultivated in the presence of discomycin A, DMSO (negative control) or vancomycin (positive control), monitoring the [M-2H]^2-^ ion of UDP-MurNAc-pentapeptide (m/z=573.6685) in ESI negative mode.

## Conclusion

Here we present DiscERN, a multi-modal bioinformatics tool that solves a critical challenge in modern genome mining: the targeted, hypothesis-driven expansion of specific natural product families. By integrating four complementary algorithms, DiscERN allows users to strategically balance discovery sensitivity with predictive precision, creating a direct and reliable path from genomic data to a prioritised list of high-confidence candidate BGCs. This tuneable approach serves diverse research goals, from exhaustive screening where no potential hit can be missed, to high-precision workflows where minimising downstream analysis and experimental effort is paramount.

We have validated this entire pipeline not only through comprehensive computational benchmarking, but also by prediction-led discovery of the novel antibiotic discomycin A from a silent BGC. The journey from an in silico prediction to an isolated and structurally characterised molecule provides a concrete demonstration of DiscERN’s real-world utility. This work therefore demonstrates both a powerful and accessible new tool for the natural products community and a clear workflow for how integrated computational and synthetic biology approaches can effectively translate the promise of the genomics era into new chemical entities.

## Supporting information

BGC analysis summary

## Author Contributions

Conceptualisation of the project (JGO), software development (JGO), data analysis (JGO, HEL), isolation of discomycin A (JGO), discomycin BGC capture (HEL), construction and testing of discomycin expression strains (HEL), bacterial MIC assays (HEL, RFL), MurNac accumulation assays (HEL), mammalian cell cytotoxicity assays (VHW), structure elucidation (EFW), NMR experiments (EFW), Marfey’s analysis (EFW), LC-MS experiments (HEL, EFW, RFL), manuscript writing (JGO, HEL, EFW), manuscript editing (JGO), funding (JGO).

## Additional Files

**Additional File 1:** Supplementary Data containing additional figures and tables referred to in the manuscript as well as material and methods.

**Additional File 2:** Summary of manual analysis of BGCs for four antibiotic families

**Additional File 3:** Zipped folder containing antiSMASH outputs for all putative hits from screen of 3,561 Actinomycetes genomes with four antibiotic families. https://zenodo.org/records/16791517

## Code Availability

Instructions and source code for installation of DiscERN: https://github.com/MaxMeta/DiscERN

