## Supplementary Data and Methods for "DiscERN: An Automated Genome Mining Tool for the Discovery of Evolutionarily Related Natural Products"

### Supplementary Methods:

#### Supplementary Methods S1 – Annotation of Actinomycetes Genomes Using antiSMASH 7.0:

Genomes for antiSMASH analysis were downloaded from the NCBI Assembly database using a python script that queried for all genomes belonging to the class Actinomycetes and refined the resulting list to include only those designated as 'representative' or 'reference' genomes. Downloaded genomes were then distributed into subdirectories and annotated using a local install antiSMASH 7.0 with the 'knownclusterblast' function enabled.

#### Supplementary Methods S2 – Compilation of Reference Data For DiscERN:

Reference BGC data for DiscERN was compiled by downloading all MiBiG entries as GenBank files (Version 3.1 (October 7, 2022) and analysing these with antiSMASH 7.0.

#### Supplementary Methods S3 – Performance Evaluation of the DiscERN Ensemble:

The performance of the DiscERN ensemble was evaluated using a multi-faceted approach to robustly estimate its effectiveness in a real-world discovery scenario. We focused on two key metrics: Recall, estimated via a Leave-One-Out (LOO) cross-validation on curated datasets, and Precision, estimated by manually curating predictions from a large, uncharacterized genomic dataset. All performance metrics were calculated independently for each of the six benchmark BGC families, and the unweighted mean of these results was also calculated to provide a measure of performance on an "average" BGC family, preventing bias from families with a higher or lower number of members or hits.

**Recall Estimation via Leave-One-Out (LOO) Cross-Validation:** To estimate the sensitivity of the ensemble at different confidence thresholds, a LOO cross-validation was performed for each of the six benchmark BGC families. For a given family containing *N* members, the process was repeated *N* times. In each fold *i*, the *i*-th BGC was held out while the remaining *N-1* BGCs were used to define the family model. The held-out BGC was then classified, and the number of algorithms supporting its correct identification, $k_{hits}$, was recorded.

The recall for a given confidence threshold *k* (where $k \in \{1, 2, 3, 4\}$) was calculated as the fraction of BGCs in the family that were successfully identified by *k* or more algorithms. Let $TP_k$ be the count of true positives identified by at least *k* algorithms. The recall, $R_k$, is given by:

$$R_k = \frac{TP_k}{N} = \frac{\sum_{i=1}^{N} \mathbb{I}\left(k_{hits,i} \geq k\right)}{N}$$

where $\mathbb{I}(condition)$ is the indicator function, which is 1 if the condition is true and 0 otherwise.

**Precision Estimation and Handling of Undetermined Cases:** To estimate precision, DiscERN was applied to a large dataset of 59,236 BGCs derived from 3,561 Actinomycete genomes. For each confidence tier *k* (where hits are supported by exactly *k* algorithms), putative hits were manually curated. Owing to large numbers of hits, especially at lower confidence tiers, a maximum of 20 hits were examined for each confidence tier.

Each BGC was manually classified as a True Positive (TP), False Positive (FP), or Not Determined (ND) if fragmentation prevented a confident assignment. To calculate a final precision estimate for each tier, we first calculated the precision based only on the subset of classifiable hits. To account for the ND cases, we calculated two sets of precision estimates: a primary estimate and a conservative "worst-case" estimate.

*Let, for each tier k:*

- $N_k$ = Total number of hits predicted by DiscERN.
- $tp_k, fp_k, nd_k$ = The counts of TPs and FPs and NDs respectively.

In cases where $N_k > 20$ , the precision was estimated using 20 randomly selected hits. Otherwise the entire collection of hits was used. Our primary estimate assumes that the ND cases are distributed in the same proportion as the classifiable cases. Here, the precision for the classifiable subset, $P_{det,k}$, was first calculated:

$$P_{det,k} = \frac{tp_k}{tp_k + fp_k}$$

This precision was then applied to the entire population of $N_k$ hits for that tier to estimate the total adjusted True Positive ($\widehat{TP_k}$) and False Positive ($\widehat{FP_k}$) counts. This approach assumes that the ND cases are distributed in the same proportion as the classifiable cases.

$$\widehat{TP_k} = N_k \times P_{det,k}$$

$$\widehat{FP_k} = N_k \times (1 - P_{det,k})$$

For our conservative "worst-case" estimate, we assumed that all ND cases were False Positives. In this scenario, the total True Positives are only those directly observed and extrapolated, while False Positives include both observed FPs and all NDs. The conservative precision, $P_{cons,k}$, is thus calculated from the sample counts as:

$$P_{cons,k} = \frac{tp_k}{tp_k + fp_k + nd_k}$$

The corresponding conservative estimates for the total population were then calculated by applying this rate to the $N_k$ hits.

**Calculation of Cumulative Precision and F1 Score:** Using both sets of adjusted counts (primary and conservative), we calculated the cumulative precision ($CP_k$) for a threshold of *k* is given as:

$$CP_k = \frac{\sum_{i=k}^{4} \widehat{TP_i}}{\sum_{i=k}^{4} (\widehat{TP_i} + \widehat{FP_i})}$$

So, for example:

$$CP_3 = \frac{\widehat{TP_3} + \widehat{TP_4}}{\widehat{TP_3} + \widehat{TP_4} + \widehat{FP_3} + \widehat{FP_4}}$$

To provide a single metric summarizing the balance between precision and recall, an estimated F1 score was calculated for each cumulative threshold. The F1 score, $F1_k$, is the harmonic mean of the estimated cumulative precision ($CP_k$) and the estimated recall from the LOO analysis ($R_k$):

$$F1_k = 2 \times \frac{CP_k \times R_k}{CP_k + R_k}$$

#### Supplementary Methods S4 – BGC Cloning and Heterologous Expression

The discomycin BGC boundary was chosen by comparison to the known boundaries of the CDA BGC from Streptomyces coelicolor. CRISPR RNAs (crRNAs) for Cas12a excision of the targeted BGC flanking sequence were designed and prepared according to previous published protocol.[1] Briefly, crRNA was synthesised using HiScribe T7 Quick High-Yield RNA Synthesis Kit (NEB) from an annealed dsDNA oligo template with unique spacer sequence targeting either upstream or downstream of the BGC. The BGC was then cloned from purified genomic DNA using Cas12a directed capture cloning.

CRISPR-Cas12a-mediated direct capture method [1] was used with modification. Briefly, the blue-white screening *lacZ* gene in pBE45 vector was replaced with PheS negative selection marker with strong constitutive promoter J23100 and pET-RBS. This reduces empty background colonies during cloning stage since empty vector colonies would express the PheS gene and exerts negative selection pressure in presence of suicide substrate 4-chlorophenylalanine. N-terminal His6-FnCas12a was purified from IPTG-induced BL21 DE3 *E. coli* cells harbouring the expression construct pET-FnCas12a-TEV (gifted by Huimin Zhao) using HisTrap column on a AKTA Pure FPLC system and stored as 10 µM aliquots in storage buffer (20 mM Tris-HCl pH 8, 150 mM NaCl, 15% (v/v) glycerol) at −80 ºC. The genomic DNA was prepared from *S. kanamyceticus* ATCC12853 using the salting out method[2] and digested using Cas12a guided by crRNA targeting BGC flanking sequences.

PCR fragments amplified from pBE48 and modified pBE45 (pBE45-*pheS*) containing homologous arms to the *dsc* cluster were mixed with digested gDNA for ligation by T4 DNA exo + fill-in protocol as described previously.[1] The ligation mixture was desalted then transformed into electrocompetent NEB10b containing the pBE14 helper plasmid and plated on LB agar with 100 µg/mL apramycin, 0.2 % D-glucose and 2.5 mM 4-chlorophenylalanine. Colonies were checked using BGC-specific primers to verify presence of BGC, and cloned BGC plasmids were purified using NucleoBond Xtra Midi kit (Macherey-Nagel) and verified by KpnI-HF (NEB) restriction digest. Verified BGC plasmid was conjugated into *S. albus* Del14 via ET12567/pUZ8002 *E. coli* conjugation donor strain, generating Del14::dsc.

Exconjugants were recovered on fresh ISP4 (Himedia laboratories), re-streaked onto ISP4 plates containing, 50 µg/mL apramycin and 50 µg/mL nalidixic acid, and verifying the presence of the integrated *dsc* cluster by PCR. Spore stocks were made and stored at −80 ˚C in 10 % glycerol. All PCR cloning steps used Q5 High-Fidelity 2X Master Mix (NEB). For colony screening BioMix Red 2X reaction mix (Bioline) was used. All oligonucleotides used are listed in Table S1. Antibiotic selection was performed using the following concentrations: kanamycin 50 µg/mL, apramycin 50 µg/mL, spectinomycin 50 µg/mL, nalidixic acid 50 µg/mL, ampicillin 100 µg/mL.

##### Cloning of SARP-activation plasmid and heterologous expression

SARP gene (*dscF*) associated with *dsc* BGC was amplified from *S. kanamyceticus* genomic DNA with Q5 HiFi DNA Polymerase (NEB) and cloned downstream of a strong constitutive promoter A21RS into pOJ436 (with PhiC31 integrase system) or pIJ10257 (with PhiBT1 integrase system) with NEBuilder HiFi Assembly Master mix. Cloned plasmids were verified with PCR and Sanger sequencing before conjugating into *S. kanamyceticus* (PhiC31) or Del14*::dsc*(PhiBT1) via ET12567 *E. coli* conjugation donor strain, generating Skan::dsc*F* and Del14::*dsc-dscF* respectively. Exconjugants were recovered from fresh ISP4 re-streaked plates supplemented with appropriate antibiotics (PhiBT1: 100 ug/mL hygromycin; PhiC31: 50 ug/mL apramycin; both: 50 ug/mL nalidixic acid) and checked with PCR to verify BGC integration, and spore stocks were harvested and stored at -80 oC.

#### Supplementary Methods S5 – Isolation of Discomycin A:

A single colony colonies of Del14::*dsc-dscF* (PhiBT1) from 5-day old ISP4 agar was inoculated into 50 mL autoclaved tryptic soy broth (TSB) media and incubated at 30 ˚C 200 rpm for 2 days. Pre-culture was visually inspected for contamination before inoculating 0.5% (v/v) into 10 x 1L liquid TSB medium in baffled 2.5 L culture flasks containing 25 g/flask sterile HP-20 resin incubated Fermentation was allowed to proceed at 30 ˚C for 10 days before collection of HP-20 resin by filtration. HP-20 was washed extensively with distilled water and packed into a glass column. The column was then sequentially eluted with 500 mL each of 20, 40, 60, 80 and 100 % acetonitrile. Fractions were then checked for calcium-dependent antibiotic activity, which was found predominantly in the 60 % fraction. This was evaporated to dryness under reduced pressure and redissolved in 15 mL of aqueous 0.1 % (v/v) triethylamine. Insoluble compounds were removed by centrifugation, before addition of $CaCl_2$ to a final concentration of 100 mM to induce precipitation of discomycin. After a 20 min incubation on ice, The precipitated discomycins were collected by filtration and the aqueous phase decanted. The precipitated discomycins were then washed using acetonitrile to remove lipids before being dissolved in a minimal volume of methanol. Semi-preparative HPLC was performed to separate the discomycins using a VP 150/10 Nucleodur C-18, HTEC, 5 µm column (Macherey-Nagel, Germany) on a 1260 Infinity II HPLC (Agilent, USA).

#### Supplementary Methods S6 – Marfey's Analysis of Discomycin A:

*N*α-(2,4-Dinitro-5- fluorophenyl)-L-alaninamide (L-FDAA) and *N*α-(2,4-dinitro-5-fluorophenyl)-D-alaninamide (D-FDAA) were purchased from Santa Cruz Biotechnology. Amino acids and standards were purchased from Chem-impex int'l Inc, Sigma, or Merck. HPLC-grade MeCN and MeOH were degassed and filtered through a 0.45 µm polytetrafluoroethylene (PTFE) membrane prior to use. Water for HPLC was filtered through High-Q 103S Still. All other chemicals and solvents were of analytical grade. A sample of discomycin A (500 µg) in 6 M HCl (500 µL) was heated to 100 °C in a sealed vial for 48 h (monitored by mass spec), after which the hydrolysate was concentrated to dryness via reduced pressure. The hydrolysate was then treated with 1 M NaHCO3 (100 µL) and the sample was split into two even fractions. To each L-FDAA or D-FDAA (1 % solution in acetone, 500 µL) was added and heated at 40 °C for 1 h, after which the reaction was neutralised using 1M HCl (50 µL), diluted with ACN (500 µL) and filtered (0.45 µm PTFE) prior to HPLC-DAD-ESIMS analysis. An aliquot (10 µL) of each analyte was injected into an Agilent Zorbax SB-C3 column, 5 µm, 150 × 4.6 mm, 50 °C, with a 1 mL/min, 55 min linear gradient elution from 15 % to 60 % MeOH/$H_2O$ with a 5 % isocratic modifier of 1 % formic acid in MeCN (5-minute hold at 60 %, 5-minute re-equilibration at 15%). The analyte amino acid content was assessed by UV (340 nm) and ESI(+)MS monitoring, compared to authentic standards. D/L-Amino acids (L-Ser, L-Thr, L-Allo-Thr, L-Asp, D-Hpg, Gly, L-Asn, L-Ilu, L-Allo-Ilu) standards (50 µl of 50µM solutions) were dissolved in 20 µL of 1 M $NaHCO_3$. This was done in duplicate, and to each L-FDAA or D-FDAA (1 % solution in acetone, 100 µL) was added and heated at 40 °C for 1 h, after which the reaction was neutralised with 1 M HCl (20 µL), diluted with MeCN (500 µL) and filtered (0.45 µm PTFE) prior to HPLC-DAD-ESIMS analysis as above. In order to obtain a reference retention time for β-Me-Glu, which was not available for purchase, daptomycin was hydrolysed and analysed as described above.

#### Supplementary Methods S7 – Elucidation of the Structure of Discomycin A:

The structure of discomycin A was elucidated using a combination of high-resolution electrospray ionisation mass spectrometry (HRESI-MS), 1D and 2D NMR experiments, and C3 Marfey's analysis.[3] NMR data was acquired in deuterated methanol at 25 °C on a JEOL JNM-ECZ600R spectrometer equipped with a 5 mm FG/RO digital autotune probe (600 MHz for $^1$H nuclei and 150 MHz for $^{13}$C nuclei). The residual solvent peak was used as an internal standard (δC 49.00; δH 3.31). An Agilent 6530 Q-TOF coupled to an Agilent 1260 HPLC system was used to obtain HRESI-MS data. The observed mass of discomycin A (*m/z* 1415.5865 $[M + H]^+$) corresponds to a molecular formula of $C_{62}H_{87}N_{12}O_{26}$ (calcd, 1415.5849; Δ =1.13 ppm) (**Figure S2**).

Analysis of the NMR data (COSY, HSQC, HMBC) showed the presence of spin systems attributable to three aspartic acids (Asp), asparagine (Asn), serine (Ser), threonine (Thr), glycine (Gly), and isoleucine (Ilu), β-hydroxyphenylalanine (β-OH-Phe), 4-hydroxylphenylglycine (Hpg) and β-methylglutamate (β-Me-Glu), (**Figure S4**). The N-terminal acyl substituent was determined to be 2-hydroxyoctanoic acid, consisting of one carbonyl (δc 177.4), one oxymethine (δc 72.9), five methylenes (δc 35.6, 32.9, 30.2, 26.1, 23.7, 14.4) and one methyl (δc 14.4). Elucidation of the lipid tail was achieved via COSY correlations between H-2 (δh 4.07), $H_2$-3 (δh 1.77, 1.61), $H_2$-4 (δh 1.45, 1.43), $H_2$-5(δh 1.30), along with connections between $H_2$-7 (δh 1.31) and $H_3$-8 (δh 0.89). The overlapping protons of $H_2$-5, $H_2$-6, $H_2$-7 (δh 1.30-1.31) obscured COSY correlations, but connection of these carbons could be established via HMBC correlations from $H_3$-8 to C-7 (δc 23.7) and C-6 (δc 32.9) and $H_2$-4 to C-5 (δc 30.2) and C-6.

The configuration of chiral centres in each amino acid was defuced using $C_3$ Marfey's analysis(**Figure S7, Table S6**). HMBC correlations from each amino acid alpha protons to the nearby neighbouring carbonyl defined connectivity of the molecule, with the exception $Ilu_{11}$ where HMBC correlations from the beta carbon protons established connectivity (Figure S5). The connectivity was corroborated by ESI-MS/MS fragmentation (Figure S6)

#### Supplementary Methods S8 – Broth Dilution MIC Assays:

All MIC bioassays and mechanism of action studies were performed in LB broth supplemented with various concentraqtions of $CaCl_2$.A single colony of indicator strain (*Bacillus subtilis* E168, *Staphylococcus aureus* ATCC25923, methicillin-resistant *Staphylococcus aureus* USA 300, *Acinobacter baumannii* Ab5075, *Escherichia coli* ΔtolC) was resuspended in 1 mL media and the OD600 was measured. Bacteria suspensions were diluted to OD600 = 0.0005 with media and kept aside while two-fold drug dilution plates were prepared in a final volume of 100 µL MHB medium supplemented with $CaCl_2$. 50 µL diluted culture of indicator strains were added to corresponding wells. Plates were sealed with sterile breathable membrane, wrapped in aluminium foil, and incubated at 37 ºC statically (without shaking) for 18 h. The lowest concentrations of antibiotic that resulted in no visible growth of indicator strains were recorded as the MIC. The broth dilution MIC bioassays were carried out in biological triplicate (n=3).

#### Supplementary Methods S9 – Mammalian Cytotoxicity Assay:

Cytotoxicity was assessed against the human carcinoma cell line, HCT116, using a standard 48 hour cell proliferation assay. In brief, cells were plated into the wells of a 96-well plate at a seeding density of 1.2 x $10^4$ cells/well before incubating for 24 hours (37°C, 5% $CO_2$). Cells were treated with discomycin A at various concentrations then incubated for a further 48 hours (n = 2–3 independent experiments each comprising of two technical replicates). MTS was added and the plate was incubated for 1–2 hours, then the absorbance measured at 480 nm on a spectrophotometer. T

**Supplementary Methods S10 – LC-MS/MS Analysis:**

Crude extracts and purified compounds were dissolved in LC-MS grade MeOH, and filtered with 0.22 µm PTFE syringe filter. Each sample was analysed on an Agilent 1260 Infinity binary pump fitted with Agilent InfinityLab Poroshell 120 Reversed-Phase EC-C18 column, 100 x 2.1 mm, 2.7 µm particle size. Mobile phase A was 99.9 % $H_2O$/0.1 % $NH_4HCO_2$. Mobile phase B was 99.9 % MeCN/0.1 % HCOOH. The flow rate was 0.4 mL/min with the gradient 0–1 min 5.0% B, 1–20 min 5.0 %–100 % B, 20–23 min 100 % B, 23–28 min 5% B. MS data was collected using an Agilent 6530 Accurate Mass Q–TOF LC/MS. Data was collected in both ESI positive and negative modes with m/z range of 100–3000. The MS source settings used were: capillary voltage of 3500 V, nebulizer gas ($N_2$) pressure of 30 psig, ion source temperature of 275 °C, sheath gas temperature of 300 °C and flow of 0.4 mL/min, and the acquisition rate was three spectra/s. The ion isolation width was set wide at 7 amu. Minutes 0–1.5 were sent to waste and minutes 1.5–25 recorded. For untargeted ion fragmentation (auto–MS/MS), collision induced dissociation (CID) energies were dependent on the precursor mass determined by the pre–set equations CID = 2.62$x$ + 14.75 (low energy) and CID = 3.93$x$ + 22.13 (high energy). The four most intense ions per MS scan were subjected to CID and were actively excluded after three spectra for 0.4 min. For targeted fragmentation of purified compounds were directly injected and optimal CID energy were determined empirically.

#### Supplementary Figures and Tables:

##### Table S1 – MiBiG Accessions and Compound Names for BGCs Used to Define Antibiotic Families

| | |
|---|---|
| **CDAs** | |
| **BGC0000315** | CDA1b, CDA2a, CDA2b, CDA3a, CDA3b, CDA4a, CDA4b |
| **BGC0001370** | lipopeptide 8D1-1, lipopeptide 8D1-2 |
| **BGC0001968** | cadaside A, cadaside B |
| **BGC0001984** | sarpeptin A, sarpeptin B |
| **BGC0000291** | A54145 |
| **BGC0000336** | daptomycin |
| **BGC0000379** | glycinocin A |
| **BGC0000354** | friulimicin A, friulimicin B, friulimicin C, friulimicin D |
| **BGC0001448** | malacidin A, malacidin B |
| **BGC0002430** | gausemycin A, gausemycin B |
| **BGC0000439** | taromycin A |
| **Glycopeptides** | |
| **BGC0000289** | A40926 |
| **BGC0000290** | A-47934 |
| **BGC0000311** | balhimycin |
| **BGC0000322** | chloroeremomycin |
| **BGC0000326** | isocomplestatin |
| **BGC0000418** | Ristocetin A, Desmethyl ristocetin A, Ristocetin B, Desmethyl ristocetin B |
| **BGC0000419** | ristomycin A |
| **BGC0000440** | Teicoplanin A2-1, Teicoplanin A2-2, Teicoplanin A2-3, Teicoplanin A2-4, Teicoplanin A2-5 |
| **BGC0000441** | teicoplanin |
| **BGC0000455** | vancomycin |
| **BGC0001178** | UK-68,597 |
| **BGC0001459** | decaplanin |
| **BGC0001460** | decaplanin |
| **BGC0001461** | Actinoidin B |
| **BGC0001462** | avoparcin |
| **BGC0001635** | kistamicin A |
| **BGC0001955** | keratinimicin A, keratinimicin B, keratinimicin C, keratinimicin D |
| **BGC0002314** | corbomycin |
| **BGC0002344** | A50926 A, A50926 B |
| **BGC0002637** | rimomycin A, rimomycin B, rimomycin C |
| **BGC0002638** | misaugamycin A, misaugamycin B |
| **BGC0002702** | GP6738 |
| **BGC0002992** | pekiskomycin |
| **BGC0003168** | pekiskomycin |
| Macrolides | |
| **BGC0000054** | erythromycin |
| **BGC0000055** | erythromycin A, erythromycin B, erythromycin C, erythromycin D |
| **BGC0000085** | lankamycin |
| **BGC0000092** | megalomicin A, megalomicin B, megalomicin C1, megalomicin C2 |
| **BGC0002958** | 12-ketomethymycin N-oxide |
| **BGC0001830** | rosamicin, salinipyrone A, pacificanone A |
| **BGC0002086** | rosamicin |
| **BGC0001931** | carrimycin |
| **BGC0000096** | midecamycin |
| **BGC0002452** | leucomycin |
| **BGC0000035** | chalcomycin A |

| | |
|---|---|
| **BGC0000047** | 10,11-dihydro-8-deoxy-12,13-deepoxy-12,13-dihydrochalcomycin |
| **BGC0001396** | aldgamycin J, aldgamycin K, aldgamycin P, aldgamycin E |
| **BGC0000102** | mycinamicin II |
| **BGC0001812** | tylactone |
| **BGC0002033** | spiramycin |
| **Aminoglycosides** | |
| **BGC0000702** | kanamycin |
| **BGC0000703** | kanamycin |
| **BGC0000705** | kanamycin |
| **BGC0000704** | kanamycin |
| **BGC0000721** | tobramycin |
| **BGC0000720** | tobramycin |
| **BGC0000719** | tobramycin |
| **BGC0000696** | gentamicin |
| **BGC0000689** | 2-deoxystreptamine |
| **BGC0001603** | gentamicin |
| **BGC0000714** | sisomicin |
| **BGC0000692** | apramycin |
| **BGC0000708** | lividomycin |
| **BGC0000712** | paromomycin |
| **BGC0000709** | neomycin |
| **BGC0000711** | neomycin |
| **BGC0000713** | ribostamycin |
| **BGC0000699** | hygromycin A |
| **BGC0000700** | istamycin |
| **BGC0000724** | streptomycin |
| **BGC0000690** | hydroxystreptomycin |
| **BGC0000716** | spectinomycin |
| **Rifamycin like** | |
| BGC0000136 | rifamycin |
| BGC0000137 | rifamycin |
| BGC0001287 | chaxamycin A, chaxamycin B, chaxamycin C, chaxamycin D |
| BGC0001759 | rifamorpholine A, rifamorpholine B, rifamorpholine C, rifamorpholine D, rifamorpholine E |
| BGC0001785 | streptovaricin |
| BGC0002009 | kanglemycin A, kanglemycin V1, kanglemycin V2 |
| **Polyenes** | |
| BGC0000115 | nystatin A1 |
| BGC0001709 | nystatin |
| BGC0000116 | nystatin-like Pseudonocardia polyene A1 |
| BGC0002540 | tetramycin B |
| BGC0002106 | rimocidin |
| BGC0000034 | candicidin |
| BGC0001690 | natamycin |
| BGC0002333 | lucensomycin |
| BGC0001491 | 67-121C |
| BGC0000125 | pimaricin |
| BGC0002104 | eurocidin D, eurocidin E |
| BGC0001773 | selvamicin |

#### Table S2 Leave One Out Analysis Details

| CDAs | BS | BV | BR | ST | Sum | Recall | |
|---|---|---|---|---|---|---|---|
| BGC0000315 | 0 | 0 | 1 | 1 | 2 | K=4 | 0.09090909 |
| BGC0001370 | 0 | 1 | 1 | 1 | 3 | K>=3 | 0.33333333 |
| BGC0001968 | 0 | 1 | 1 | 1 | 3 | K>=2 | 0.81818182 |
| BGC0001984 | 1 | 0 | 0 | 1 | 2 | K>=1 | 1 |
| BGC0000291 | 0 | 0 | 1 | 0 | 1 | | |
| BGC0000336 | 0 | 0 | 1 | 1 | 2 | | |
| BGC0000379 | 0 | 1 | 1 | 1 | 3 | | |
| BGC0000354 | 1 | 1 | 1 | 1 | 4 | | |
| BGC0001448 | 0 | 0 | 1 | 0 | 1 | | |
| BGC0002430 | 0 | 0 | 1 | 0 | 1 | | |
| BGC0000439 | 0 | 0 | 1 | 1 | 2 | | |
| **Glycopeptides** | | | | | | **Recall** | |
| BGC0000289 | 1 | 1 | 1 | 1 | 4 | K=4 | 0.95833333 |
| BGC0000290 | 1 | 1 | 1 | 1 | 4 | K>=3 | 1 |
| BGC0000311 | 1 | 1 | 1 | 1 | 4 | K>=2 | 1 |
| BGC0000322 | 1 | 1 | 1 | 1 | 4 | K>=1 | 1 |
| BGC0000326 | 1 | 1 | 1 | 1 | 4 | | |
| BGC0000418 | 1 | 1 | 1 | 1 | 4 | | |
| BGC0000419 | 1 | 1 | 1 | 1 | 4 | | |
| BGC0000440 | 1 | 1 | 1 | 1 | 4 | | |
| BGC0000441 | 1 | 1 | 1 | 1 | 4 | | |
| BGC0000455 | 1 | 1 | 1 | 1 | 4 | | |
| BGC0001178 | 1 | 1 | 1 | 1 | 4 | | |
| BGC0001459 | 1 | 1 | 1 | 1 | 4 | | |
| BGC0001460 | 1 | 1 | 1 | 1 | 4 | | |
| BGC0001461 | 1 | 1 | 1 | 1 | 4 | | |
| BGC0001462 | 1 | 1 | 1 | 1 | 4 | | |
| BGC0001635 | 1 | 1 | 1 | 0 | 3 | | |
| BGC0001955 | 1 | 1 | 1 | 1 | 4 | | |
| BGC0002314 | 1 | 1 | 0 | 1 | 3 | | |
| BGC0002344 | 1 | 1 | 1 | 1 | 4 | | |
| BGC0002637 | 1 | 1 | 1 | 1 | 4 | | |
| BGC0002638 | 1 | 1 | 1 | 1 | 4 | | |
| BGC0002702 | 1 | 1 | 1 | 1 | 4 | | |
| BGC0002992 | 1 | 1 | 1 | 1 | 4 | | |
| BGC0003168 | 1 | 1 | 1 | 1 | 4 | | |
| **Macrolides** | | | | | | **Recall** | |
| BGC0000054 | 0 | 0 | 1 | 1 | 2 | K=4 | 0.625 |
| BGC0000055 | 0 | 1 | 1 | 1 | 3 | K>=3 | 0.875 |
| BGC0000085 | 1 | 1 | 1 | 1 | 4 | K>=2 | 1 |
| BGC0000092 | 0 | 1 | 1 | 1 | 3 | K>=1 | 1 |
| BGC0002958 | 0 | 1 | 1 | 1 | 3 | | |
| BGC0001830 | 1 | 1 | 1 | 1 | 4 | | |
| BGC0002086 | 1 | 1 | 1 | 1 | 4 | | |
| BGC0001931 | 1 | 1 | 1 | 1 | 4 | | |
| BGC0000096 | 1 | 1 | 1 | 1 | 4 | | |
| BGC0002452 | 1 | 1 | 1 | 1 | 4 | | |
| BGC0000035 | 1 | 1 | 1 | 1 | 4 | | |
| BGC0000047 | 1 | 1 | 1 | 1 | 4 | | |
| BGC0001396 | 1 | 1 | 0 | 1 | 3 | | |
| BGC0000102 | 0 | 1 | 1 | 0 | 2 | | |
| BGC0001812 | 1 | 1 | 1 | 1 | 4 | | |
| BGC0002033 | 1 | 1 | 1 | 1 | 4 | | |
| **Amino Glycos** | | | | | | **Recall** | |
| BGC0000702 | 1 | 1 | 1 | | 3 | K=4 | N/A |
| BGC0000703 | 1 | 1 | 1 | | 3 | K>=3 | 0.86363636 |
| BGC0000705 | 1 | 1 | 1 | | 3 | K>=2 | 0.86363636 |
| BGC0000704 | 1 | 1 | 1 | | 3 | K>=1 | 0.95454545 |
| BGC0000721 | 1 | 1 | 1 | | 3 | | |
| BGC0000720 | 1 | 1 | 1 | | 3 | | |

| | | | | | | | |
|---|---|---|---|---|---|---|---|
| BGC0000719 | 1 | 1 | 1 | | 3 | | |
| BGC0000696 | 1 | 1 | 1 | | 3 | | |
| BGC0000689 | 1 | 1 | 1 | | 3 | | |
| BGC0001603 | 1 | 1 | 1 | | 3 | | |
| BGC0000714 | 1 | 1 | 1 | | 3 | | |
| BGC0000692 | 1 | 1 | 1 | | 3 | | |
| BGC0000708 | 1 | 1 | 1 | | 3 | | |
| BGC0000712 | 1 | 1 | 1 | | 3 | | |
| BGC0000709 | 1 | 1 | 1 | | 3 | | |
| BGC0000711 | 1 | 1 | 1 | | 3 | | |
| BGC0000713 | 1 | 1 | 1 | | 3 | | |
| BGC0000699 | 1 | 1 | 1 | | 3 | | |
| BGC0000700 | 1 | 1 | 1 | | 3 | | |
| BGC0000724 | 0 | 0 | 0 | | 0 | | |
| BGC0000690 | 0 | 1 | 0 | | 1 | | |
| BGC0000716 | 0 | 0 | 1 | | 1 | | |
| **Rif** | | | | | | **Recall** | |
| BGC0000136 | 1 | 1 | 1 | 1 | 4 | K=4 | 0.83333333 |
| BGC0000137 | 1 | 1 | 1 | 1 | 4 | K>=3 | 1 |
| BGC0001287 | 1 | 1 | 1 | 1 | 4 | K>=2 | 1 |
| BGC0001759 | 1 | 1 | 1 | 0 | 3 | K>=1 | 1 |
| BGC0001785 | 0 | 1 | 1 | 1 | 3 | | |
| BGC0002009 | 1 | 1 | 1 | 1 | 4 | | |
| **Polyene** | | | | | | **Recall** | |
| BGC0000115 | 1 | 1 | 1 | 1 | 4 | K=4 | 0.83333333 |
| BGC0001709 | 1 | 1 | 1 | 1 | 4 | K>=3 | 0.91666667 |
| BGC0000116 | 1 | 1 | 1 | 1 | 4 | K>=2 | 1 |
| BGC0002540 | 1 | 1 | 1 | 1 | 4 | K>=1 | 1 |
| BGC0002106 | 1 | 1 | 1 | 1 | 4 | | |
| BGC0000034 | 1 | 1 | 1 | 1 | 4 | | |
| BGC0001690 | 1 | 1 | 1 | 1 | 4 | | |
| BGC0002333 | 1 | 1 | 0 | 0 | 2 | | |
| BGC0001491 | 1 | 1 | 1 | 1 | 4 | | |
| BGC0000125 | 1 | 1 | 1 | 1 | 4 | | |
| BGC0002104 | 1 | 1 | 1 | 1 | 4 | | |
| BGC0001773 | 0 | 1 | 1 | 1 | 3 | | |

#### Table S3 – Specificity Analysis Details

| Polymer | TP | FP | false positives per 1000 BGCs |
|---|---|---|---|
| CDA | 0 | 1 | 0.38 |
| polyene | 7 | 0 | 0.00 |
| glycopeptides | 6 | 0 | 0.00 |
| rif like | 0 | 0 | 0.00 |
| amino glycos | N/A | N/A | N/A |
| macrolides | 0 | 0 | 0.00 |
| **BS** | **TP** | **FP** | |
| CDA | 0 | 0 | 0.00 |
| polyene | 1 | 0 | 0.00 |
| glycopeptides | 5 | 1 | 0.38 |
| rif like | 0 | 0 | 0.00 |
| amino glycos | 1 | 0 | 0.00 |
| macrolides | 0 | 0 | 0.00 |
| **OL** | **TP** | **FP** | |
| CDA | 3 | 0 | 0.00 |
| polyene | 8 | 0 | 0.00 |
| glycopeptides | 6 | 0 | 0.00 |
| rif like | 0 | 0 | 0.00 |
| amino glycos | 2 | 8 | 3.06 |
| macrolides | 0 | 0 | 0.00 |
| **BV** | **TP** | **FP** | |
| CDA | 3 | 9 | 3.44 |
| polyene | 10 | 1 | 0.38 |
| glycopeptides | 6 | 6 | 2.30 |
| rif like | 0 | 0 | 0.00 |
| amino glycos | 2 | 35 | 13.39 |
| macrolides | 0 | 0 | 0.00 |

##### Figure S1. Comparison of the Discomycin and CDA BGCs

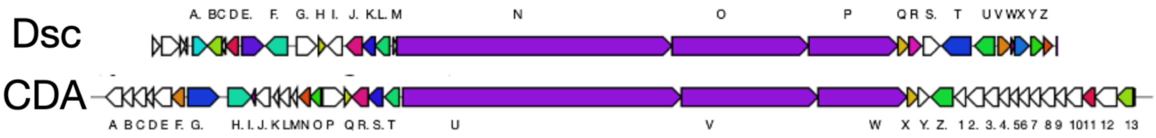

##### Table S4. Putative Functions of Genes in the Discomycin BGC

| Gene | Protein Type | Putative Function |
|---|---|---|
| **A** | prephenate dehydrogenase | Unknown |
| **B** | beta-ketoacyl-ACP synthase family protein | Fatty acid biosynthesis |
| **C** | acyl carrier protein | Fatty acid biosynthesis |
| **D** | ketoacyl-ACP synthase III | Fatty acid biosynthesis |
| **E** | fatty acyl-AMP ligase | Fatty acid biosynthesis |
| **F** | Streptomyces Antibiotic Regulatory Protein (SARP) | Regulatory |
| **G** | Histidine kinase | Regulatory |
| **H** | response regulator transcription factor | Regulatory |
| **I** | IS110 family transposase | N/A |
| **J** | PLP-dependent aminotransferase | HPG biosynthesis |
| **K** | alpha-hydroxy acid oxidase | HPG biosynthesis |
| **L** | 4-hydroxyphenylpyruvate dioxygenase | HPG biosynthesis |
| **M** | acyl carrier protein | Core Biosynthesis |
| **N** | non-ribosomal peptide synthetase | Core Biosynthesis |
| **O** | non-ribosomal peptide synthetase | Core Biosynthesis |
| **P** | non-ribosomal peptide synthetase | Core Biosynthesis |
| **Q** | Alpha-beta hydrolase | Type-II Thioesterase |
| **R** | LLM class flavin-dependent oxidoreductase | aa or fatty acid oxidation |
| **S** | M20/M25/M40 family metallo-hydrolase | Unknown |
| **Y** | HAD-IC family P-type ATPase | Transport/export |
| **U** | ABC transporter ATP-binding protein | Transport/export |
| **V** | class I SAM-dependent methyltransferase | Glutamine Methylation |
| **W** | MbtH family protein | Core Biosynthesis |
| **X** | cytochrome P450 | aa or fatty acid oxidation |
| **Y** | ABC transporter ATP-binding protein | Transport/export |
| **Z** | ABC transporter | Transport/export |

#### Figure S2. HRESI-MS Spectrum of Discomycin A

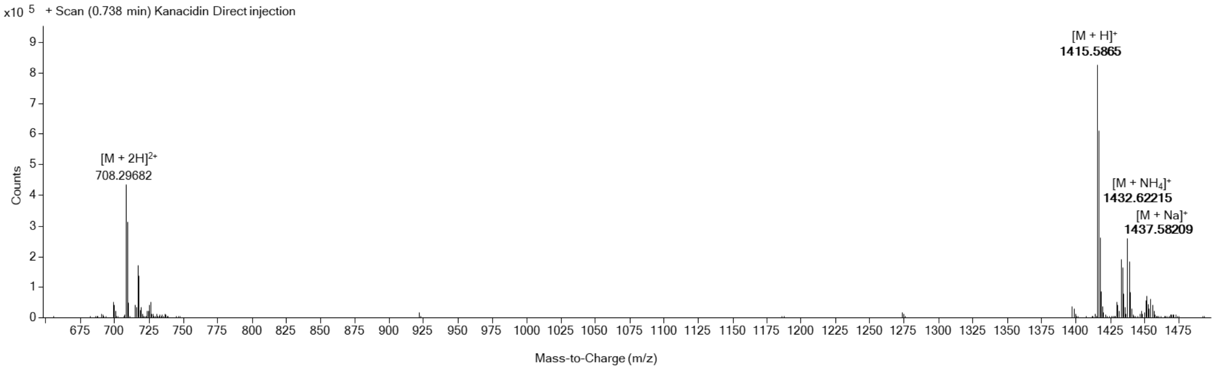

#### Figure S3. Reference Atom Numbering for Discomycin A

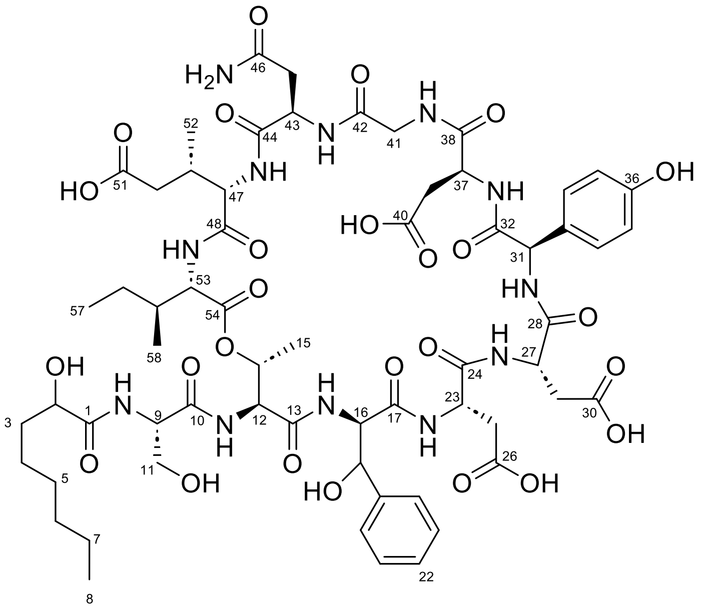

#### Table S5: Tabulated NMR Data for Discomycin A

| Pos. | $\delta_C$, type | $\delta_H$, mult. ($J$ in Hz) |
|---|---|---|
| | **2-hydroxyoctanoic acid** | |
| 1 | 177.4, C | - |
| 2 | 72.9, CH | 4.07, dd (8.0, 3.5) |
| 3 | 35.6, $CH_2$ | 1.77 |
| | | 1.61 |
| 4 | 26.1, $CH_2$ | 1.45 |
| | | 1.43 |
| 5 | 30.2, $CH_2$ | 1.30 |
| 6 | 32.9, $CH_2$ | 1.30 |
| 7 | 23.7, $CH_2$ | 1.31 |
| 8 | 14.4, $CH_3$ | 0.89 |
| | **Serine-2** | |
| 9 | 55.5, CH | 4.65 |
| 10 | 172.5, C | - |
| 11 | 63.2, $CH_2$ | 3.88, dd, 11, 5.8 |

| | | |
|---|---|---|
| | | 3.78, dd, 10.7, 6.2 |
| | **Threonine-3** | |
| 12 | 57.7, CH | 4.62, d (4.3) |
| 13 | 171.1, C | - |
| 14 | 71.0, CH | 5.26, br s |
| 15 | 16.2, $CH_3$ | 1.07 |
| | **Β-Hydroxyl phenylalanine-4** | |
| 16 | 61.4, CH | 4.66 |
| 17 | 172.5, C | - |
| 18 | 73.8, CH | 5.37, br s |
| 19 | 142.0, C | - |
| 20 | 127.4, CH | 7.38, d (7.0) |
| 21 | 129.3, CH | 7.31, t (7.0) |
| 22 | 128.7, CH | 7.25 |
| | **Aspartic Acid-5** | |
| 23 | 51.6, CH | 4.79 |
| 24 | 174.3, C | - |
| 25 | 36.4, $CH_2$ | 2.79 |
| | | 2.76 |
| 26 | 173.2, C | - |
| | **Aspartic Acid-6** | |
| 27 | 51.5, CH | 4.83 |
| 28 | 173.2, C | - |
| 29 | 36.8, CH2 | 2.93 |
| | | 2.84 |
| 30 | 174.5, C | - |

#### Figure S4. Key Correlations Defining Amino Acid Identity in Discomycin A

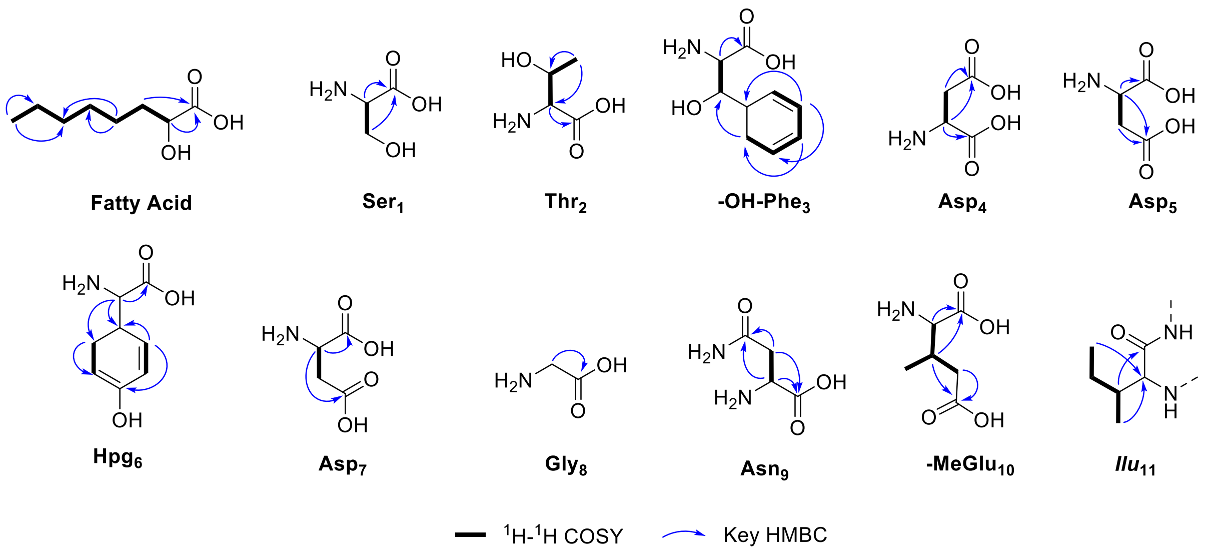

**Figure S5. Key Correlations Defining Amino Acid Connectivity in Discomycin A**

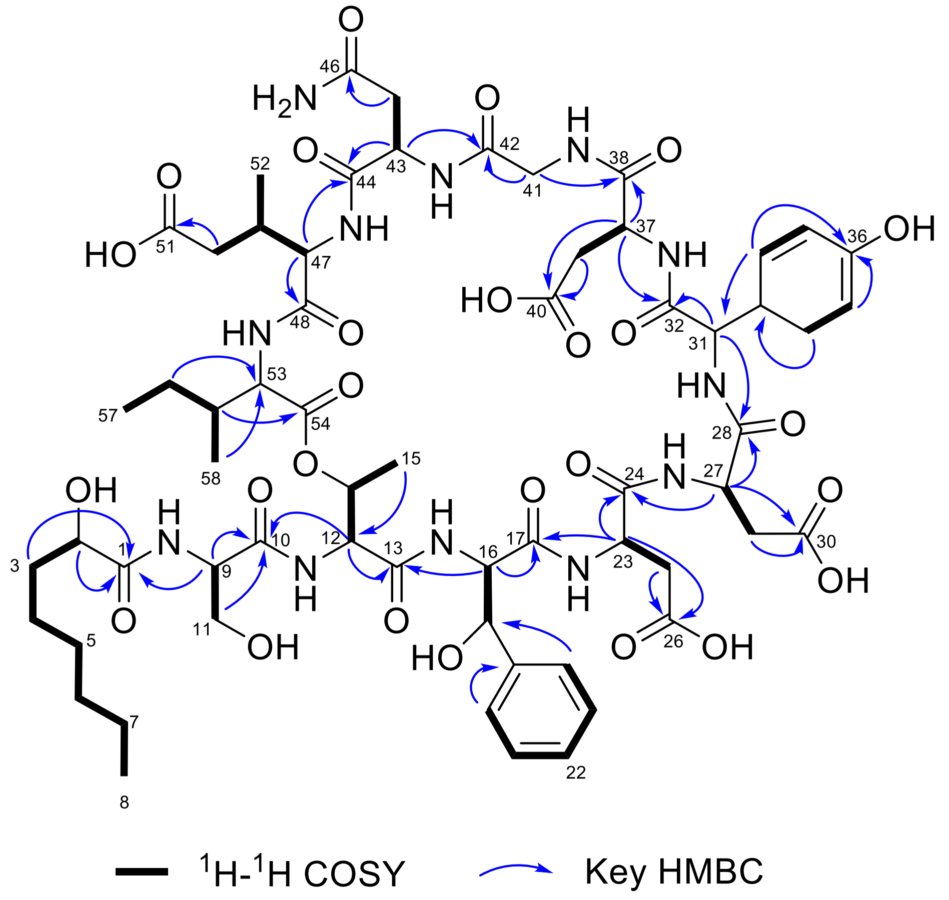

#### Figure S6. HRESI-MS/MS Fragmentation of Discomycin A

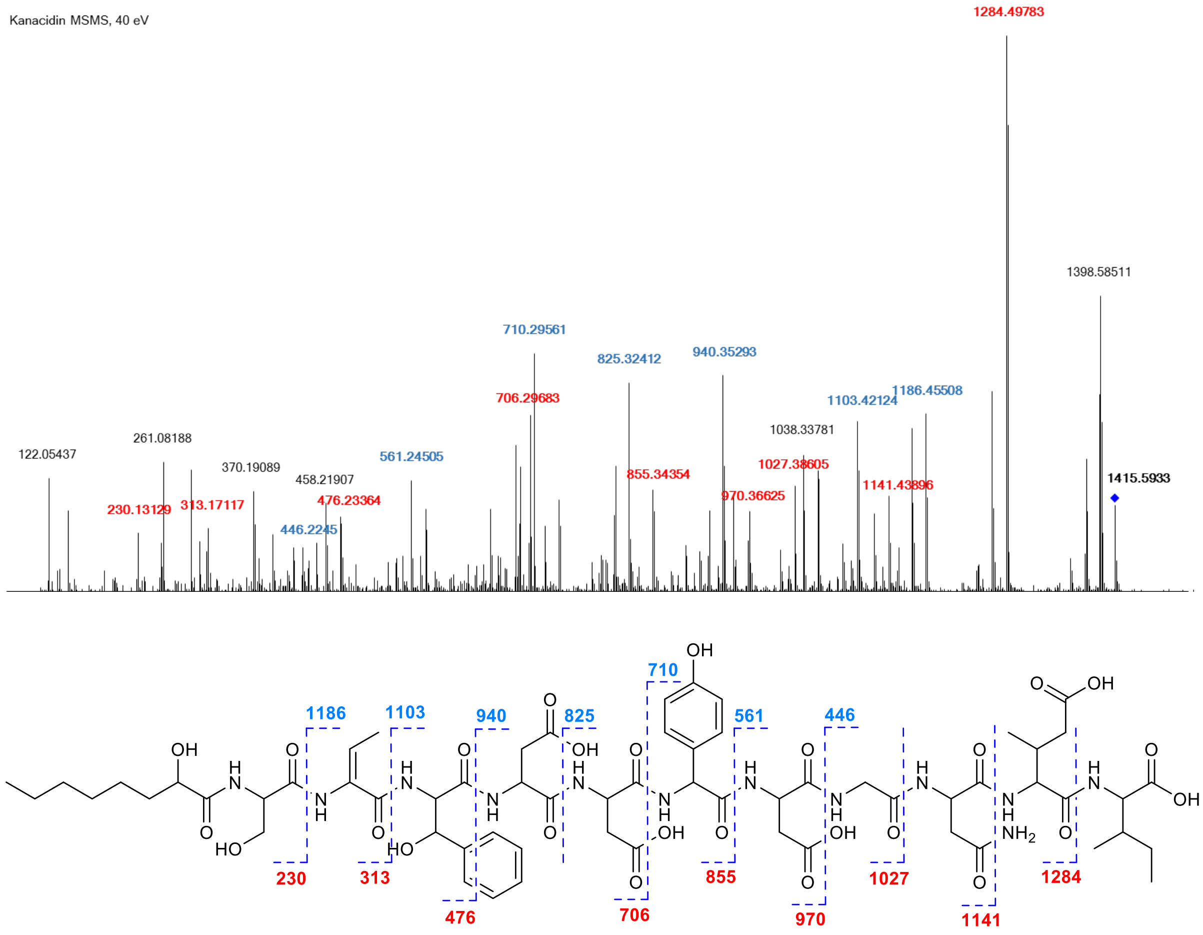

#### Table S6 – Summary of Marfey's Analysis for Discomycin A

| Amino Acid | Mass of AA + FDAA | Known Elution Order | Amino Acid standards | | Hydrolysate of discomycin | | L or D |
|---|---|---|---|---|---|---|---|
| | | | tR L-L or D-D | tR L-D or D-L | tR (AA-L-FDAA) | tR (AA-D-FDAA) | |
| Ser | 358.0993 | L→D[1] | 17.56 | 18.50 | 17.48 | 18.22 | L |
| Thr | 372.1150 | L→D[1] | 19.00 | 25.70 | 18.89 | 25.57 | L |
| Allo-Thr | 372.1150 | L→D[1] | 20.07 | 22.56 | - | - | - |
| β-OH-Phe | 434.1306 | L→D[2] | - | - | 37.88 | 30.58 | D |
| Asp | 386.0943 | L→D[1] | 22.08 | 25.40 | 22.32 | 25.4 | L |
| Hpg | 420.1150 | L→D[3] | 14.71 | 14.84 | - | - | - |
| Hpg Dimer | 672.1645 | - | 53.23 | 62.63 | 62.59 | 53.15 | D |
| Gly | 327.1150 | - | 22.67 | 22.57 | 22.65 | 22.52 | - |
| Asn | 385.1102 | L→D[1] | 17.28 | 18.15 | - | - | D* |
| β-Me-Glu | 414.1256 | L→D[4] | - | - | 27.225 | 31.33 | 2S, 3S |
| Ile | 384.1514 | L→D[1] | 39.53 | 48.05 | 39.88 | 48.44 | L |
| Allo- Ile | 384.1514 | L→D[1] | 38.39 | 47.06 | - | - | - |

[1]https://pubs.acs.org/doi/10.1021/acs.jnatprod.5b01125
[2]https://pubs.acs.org/doi/10.1021/acschembio.0c00663

**Figure S7: Extracted ion chromatograms for L-and D-FDAA derivatives of the hydrolysate of discomycin and amino acid standards.**

**Ser, *m/z* = 357.5993 – 358.5993**

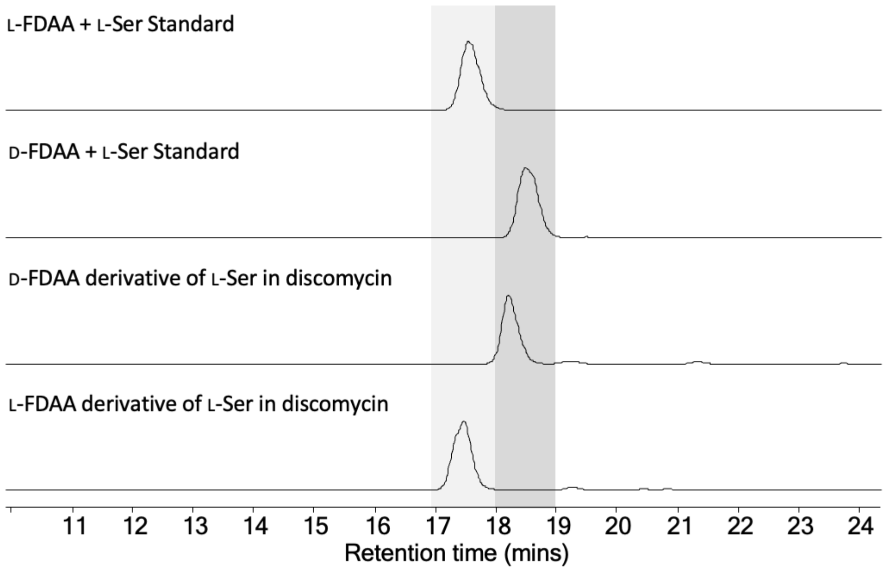

**Thr, *m/z* = 371.615 – 372.625**

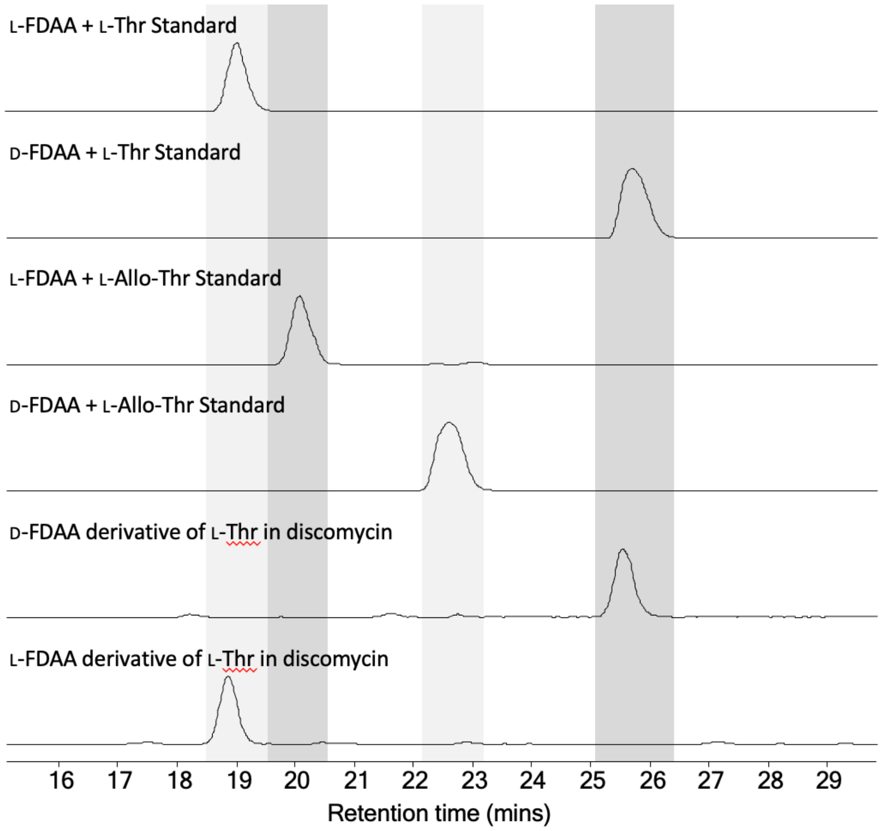

**Asp, *m/z* = 385.5943 – 386.5943**

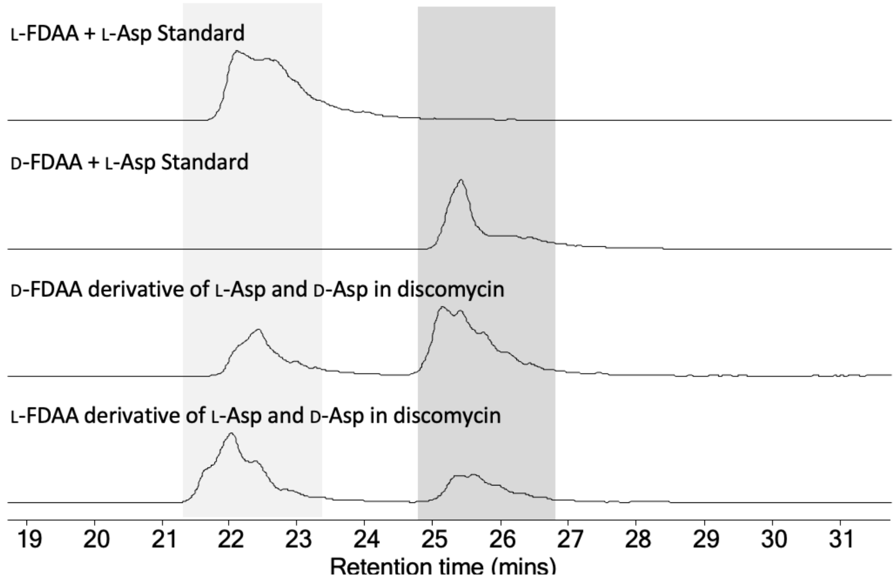

**β-OH-Phe, *m/z* = 433.6306 – 434.6306**

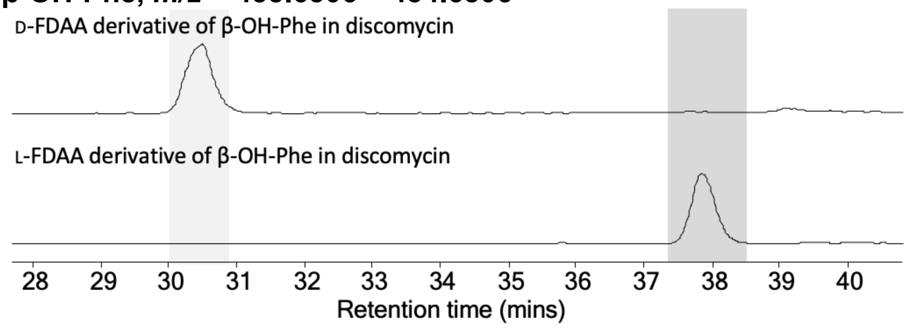

**Hpg, *m/z* = 671.6645 – 672.6645**

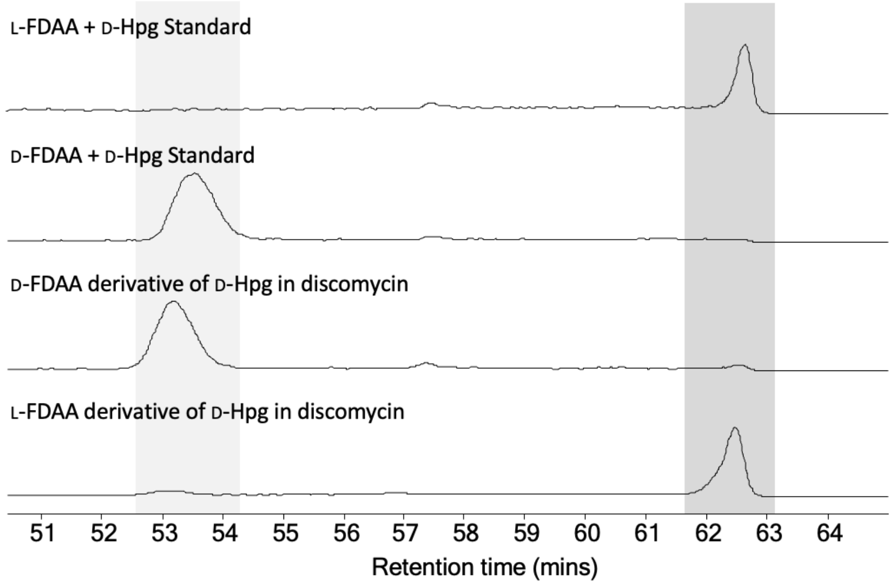

**Gly, *m/z* = 327.5888 – 328.5888**

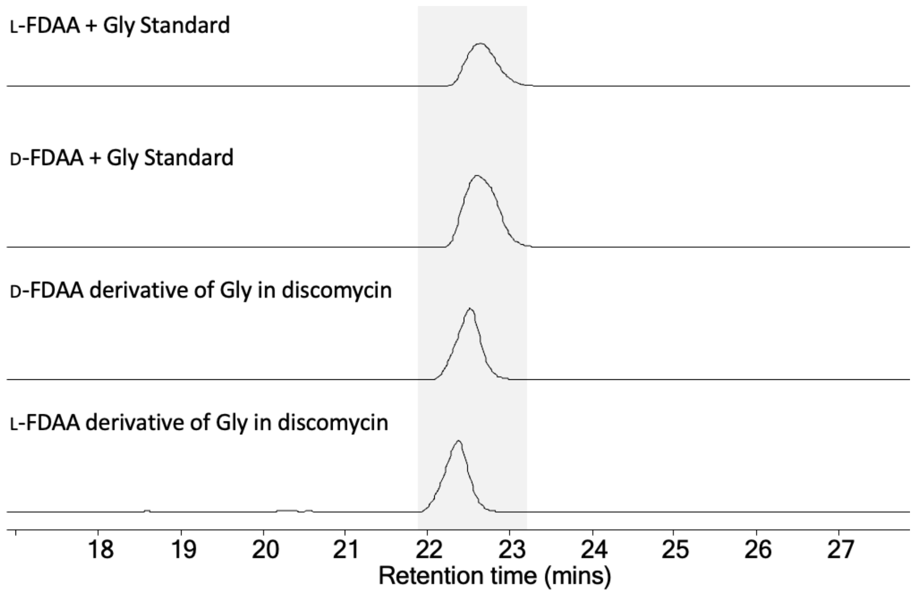

**β Me-Glu, *m/z* = 413.6256 – 414.6256**

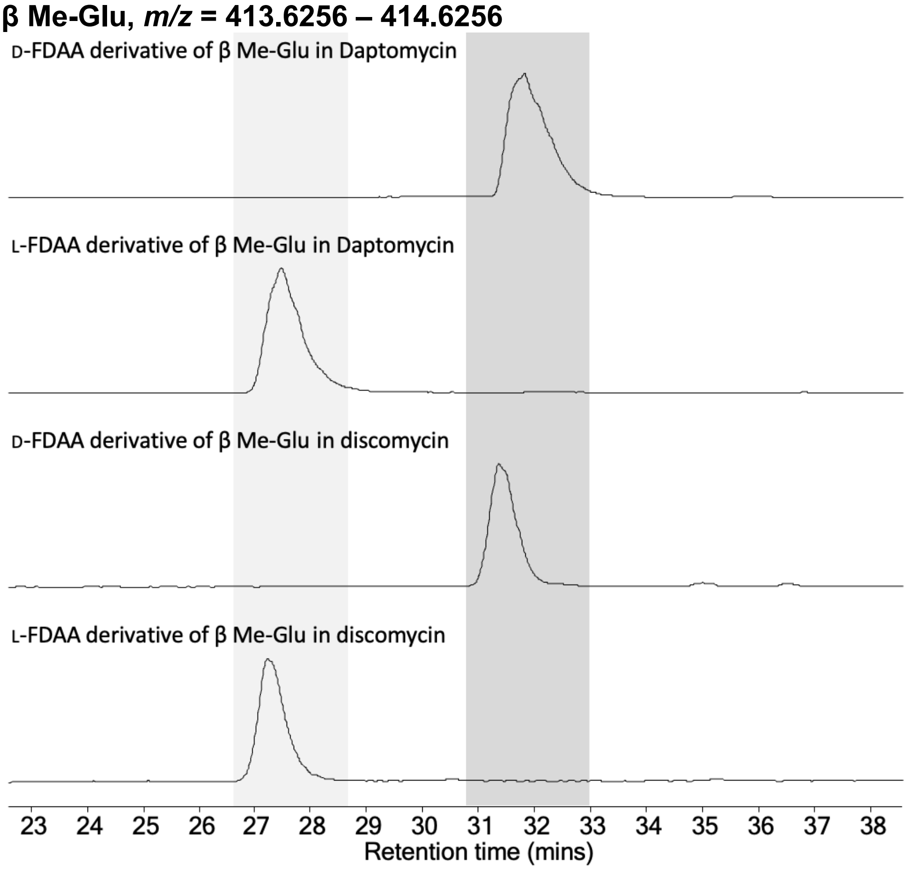

**Ile, *m/z* = 383.6514 – 384.6514**

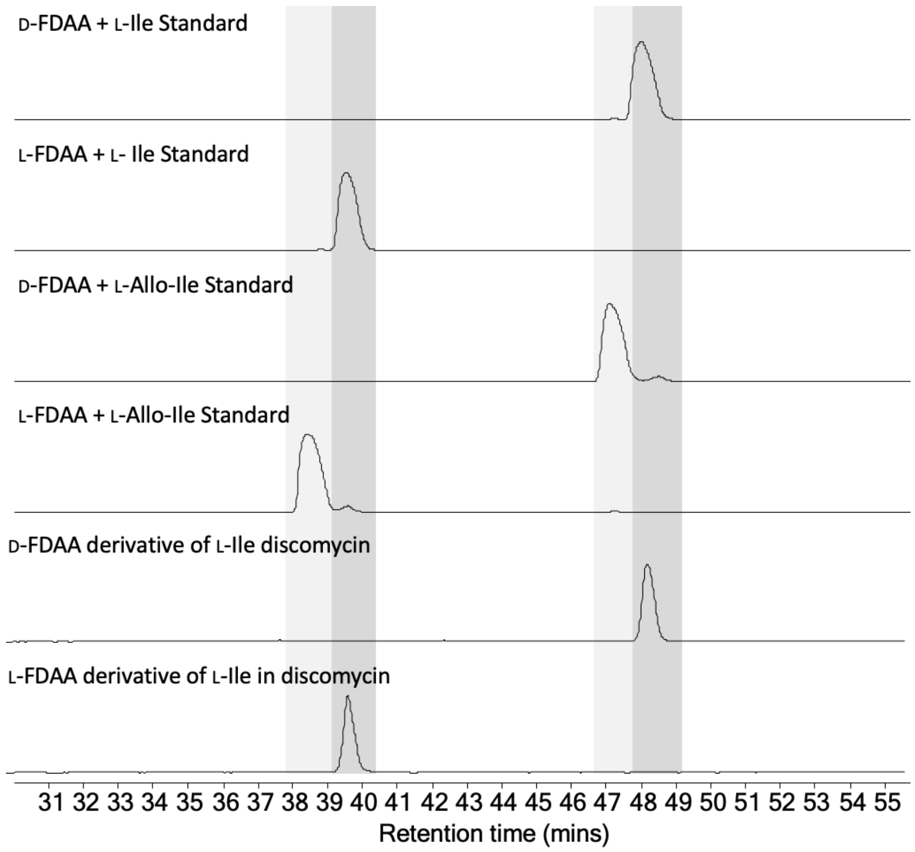

D-FDAA + L-Ile Standard
L-FDAA + L- Ile Standard
D-FDAA + L-Allo-Ile Standard
L-FDAA + L-Allo-Ile Standard
D-FDAA derivative of L-Ile discomycin
L-FDAA derivative of L-Ile in discomycin
31 32 33 34 35 36 37 38 39 40 41 42 43 44 45 46 47 48 49 50 51 52 53 54 55
Retention time (mins)

**Figure S8. 1H spectrum of discomycin A**

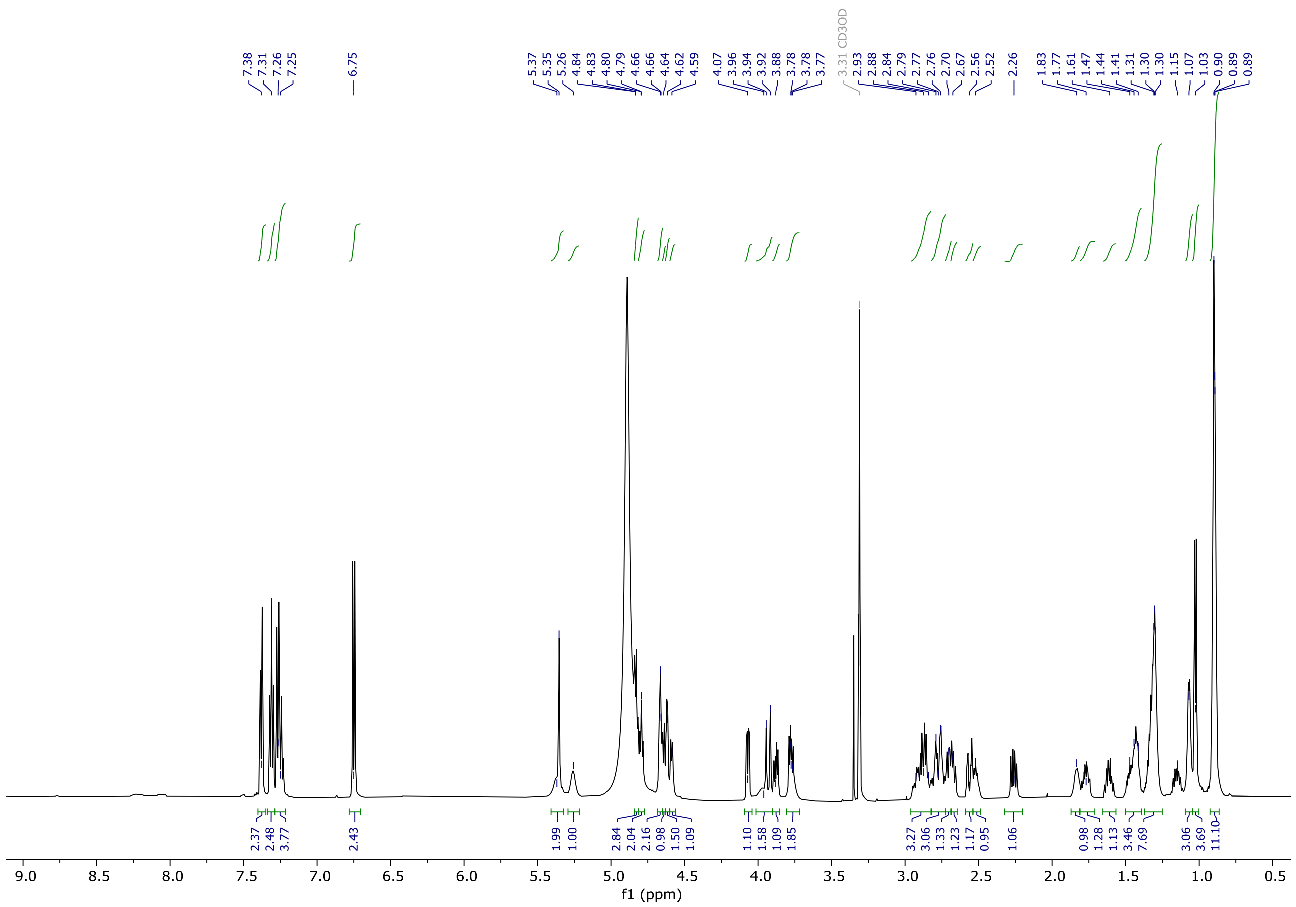

#### Figure S9. 1H-1H COSY Spectrum of discomycin A

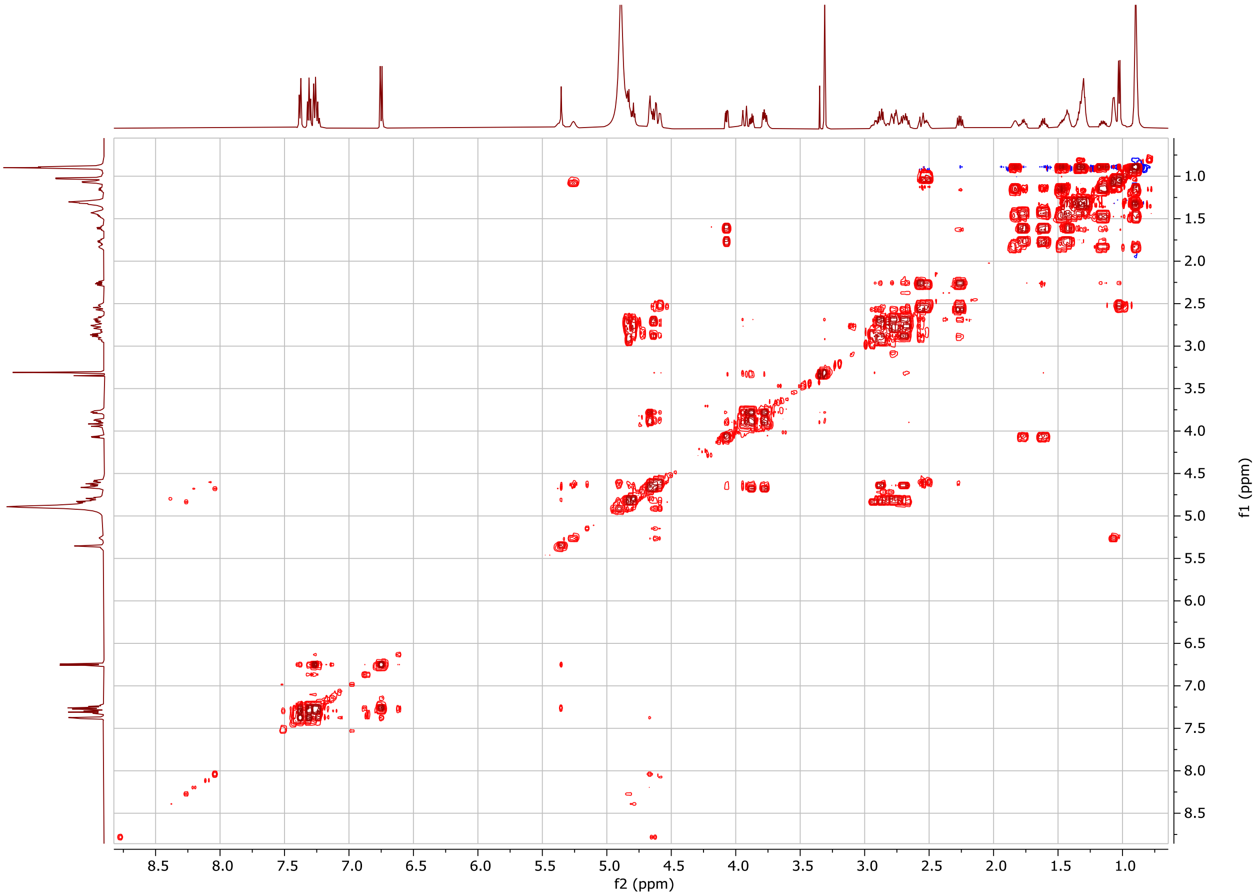

#### Figure S10. 13C spectrum of discomycin A

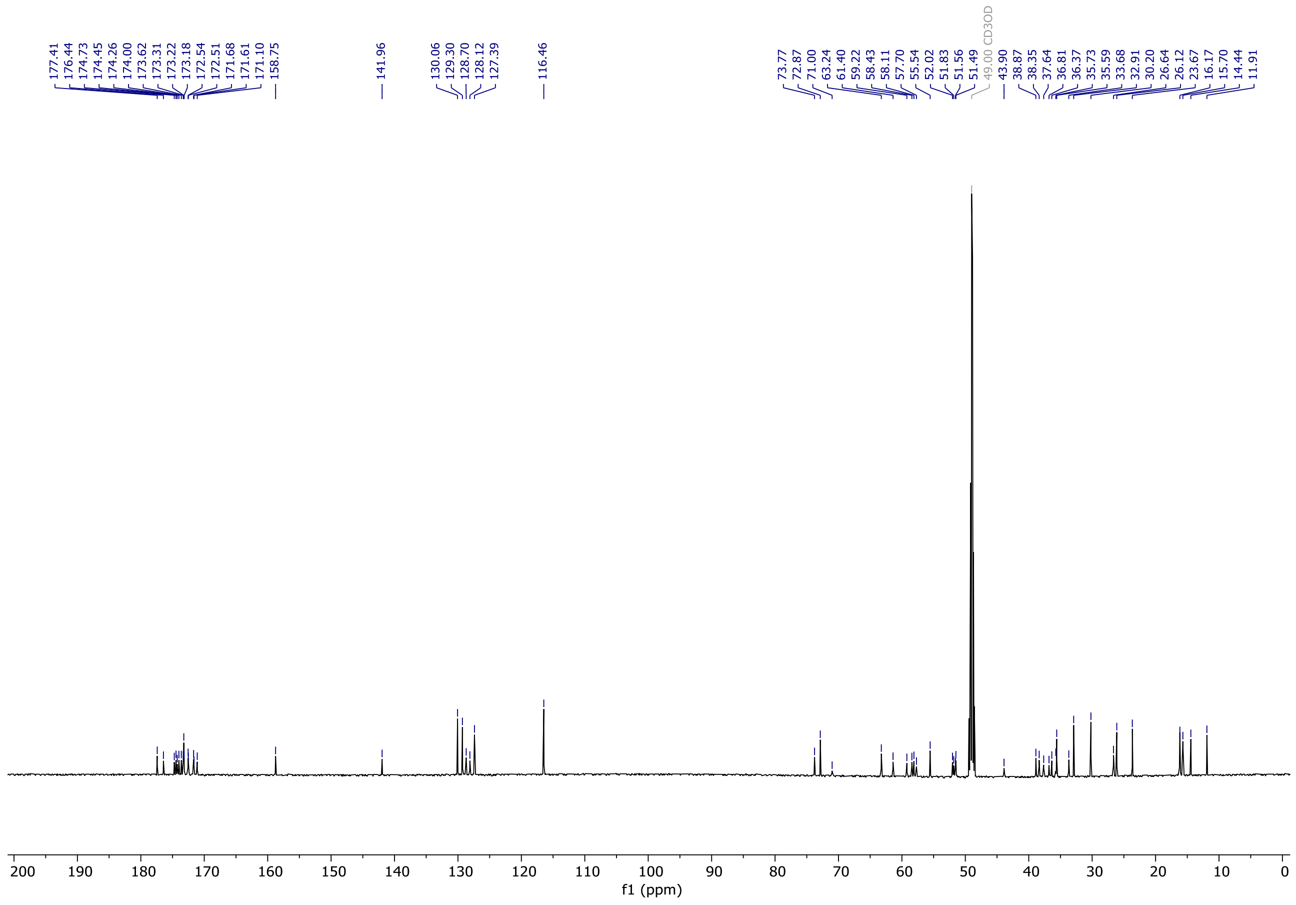

#### Figure S11. 1H-13C HSQC spectrum of discomycin A

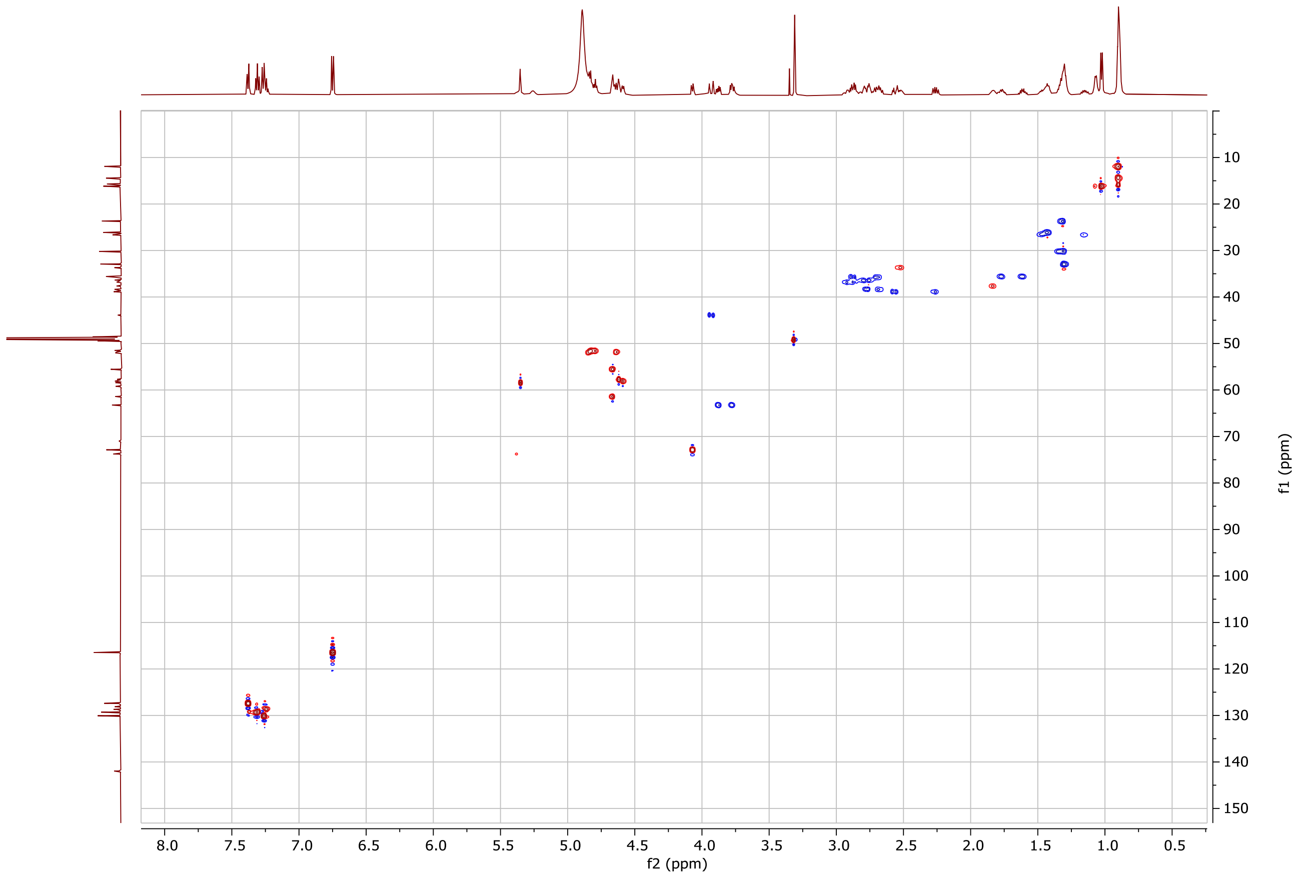

#### Figure S12. 1H-1H ROESY spectrum of discomycin A

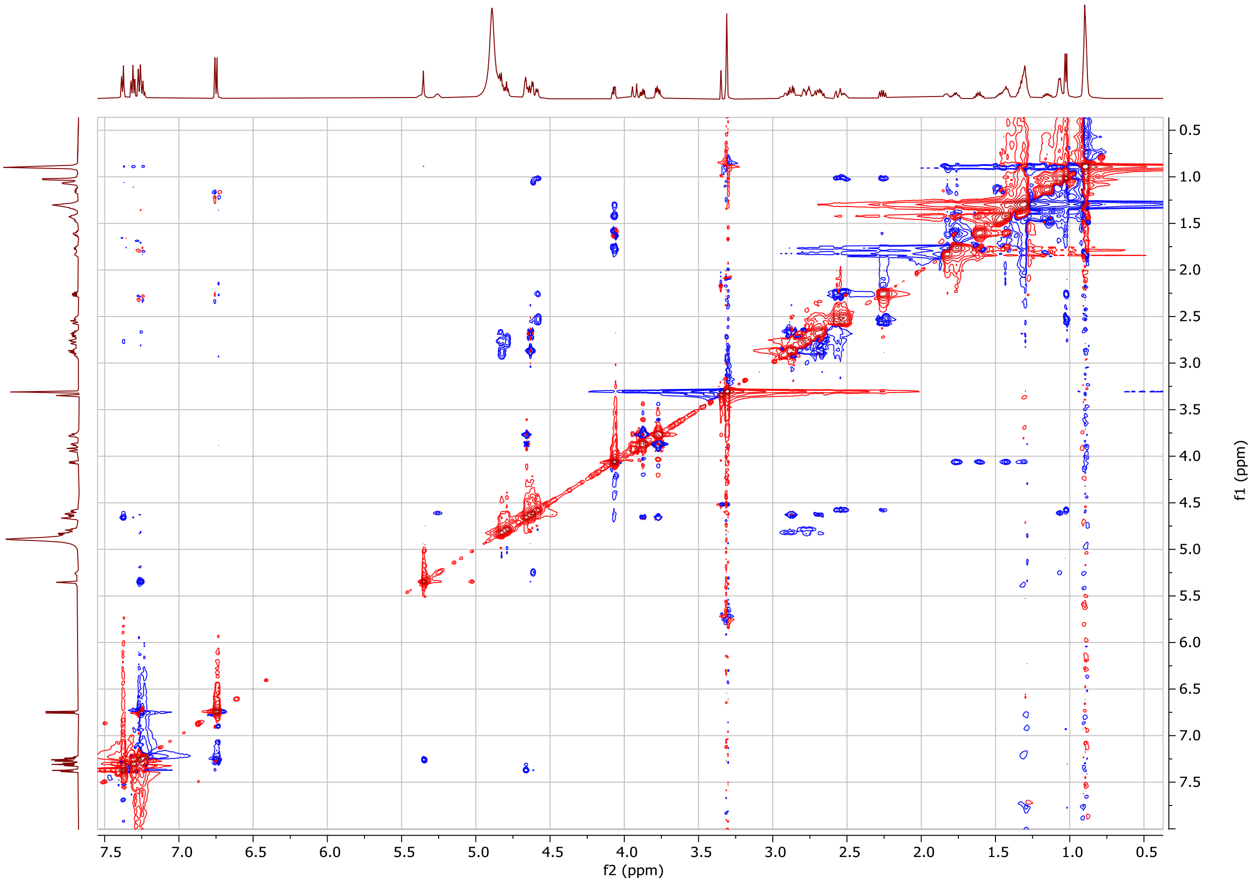

**Figure S13. 1H-13C HMBC spectrum of discomycin A**

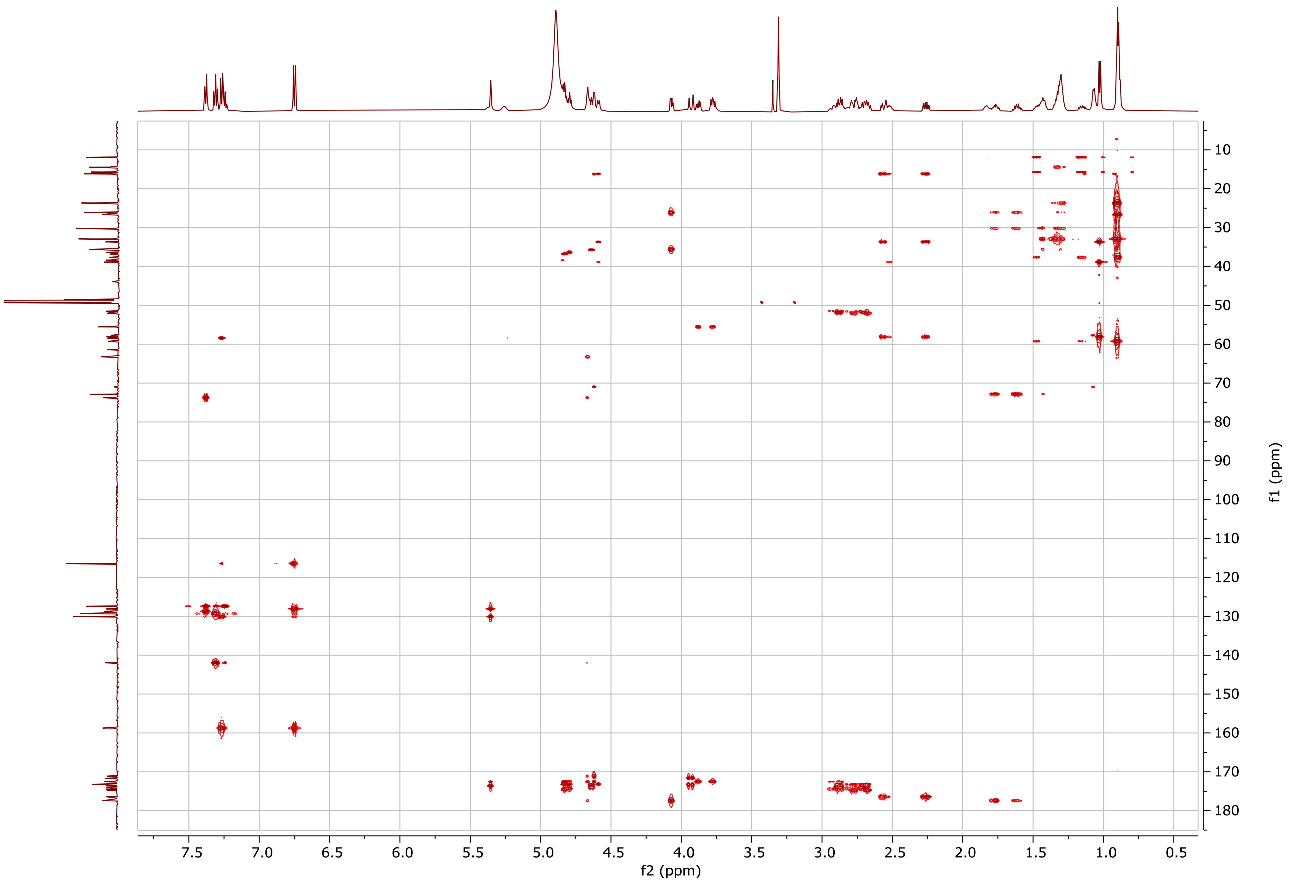

##### Table S7. MIC values for discomycin A

| Ca2+(mM) | B. subtilis | S. aureus | MRSA | HCT 116 | E. coli | A. baumanii |
|---|---|---|---|---|---|---|
| **1** | 1.0 | >32 | ND | ND | ND | ND |
| **2** | 0.5 | >32 | ND | ND | ND | ND |
| **5** | 0.125-0.25 | 32.0 | ND | ND | ND | ND |
| **10** | 0.125-0.25 | 8.0 | ND | ND | ND | ND |
| **20** | 0.1 | 2.0 | 4.0 | >32.0 | >32.0 | >32.0 |
| **50** | 0.1 | 1.0 | ND | ND | ND | ND |
| **100** | 0.031-0.063 | 0.3 | ND | ND | ND | ND |

Presented data are range of values for biological. Assays were conducted in LB medium with the indicated concentration of $CaCl_2$

##### Figure S14. DiscERN Output Dendrogram Showing Relatedness of Putative Hits

Hits supported by K≥3 algorithms are presented. With the exception of the rifamycin like dendrogram, only hits passing the automatic Pfam filter function of DiscERN are presented

**Macrolides**

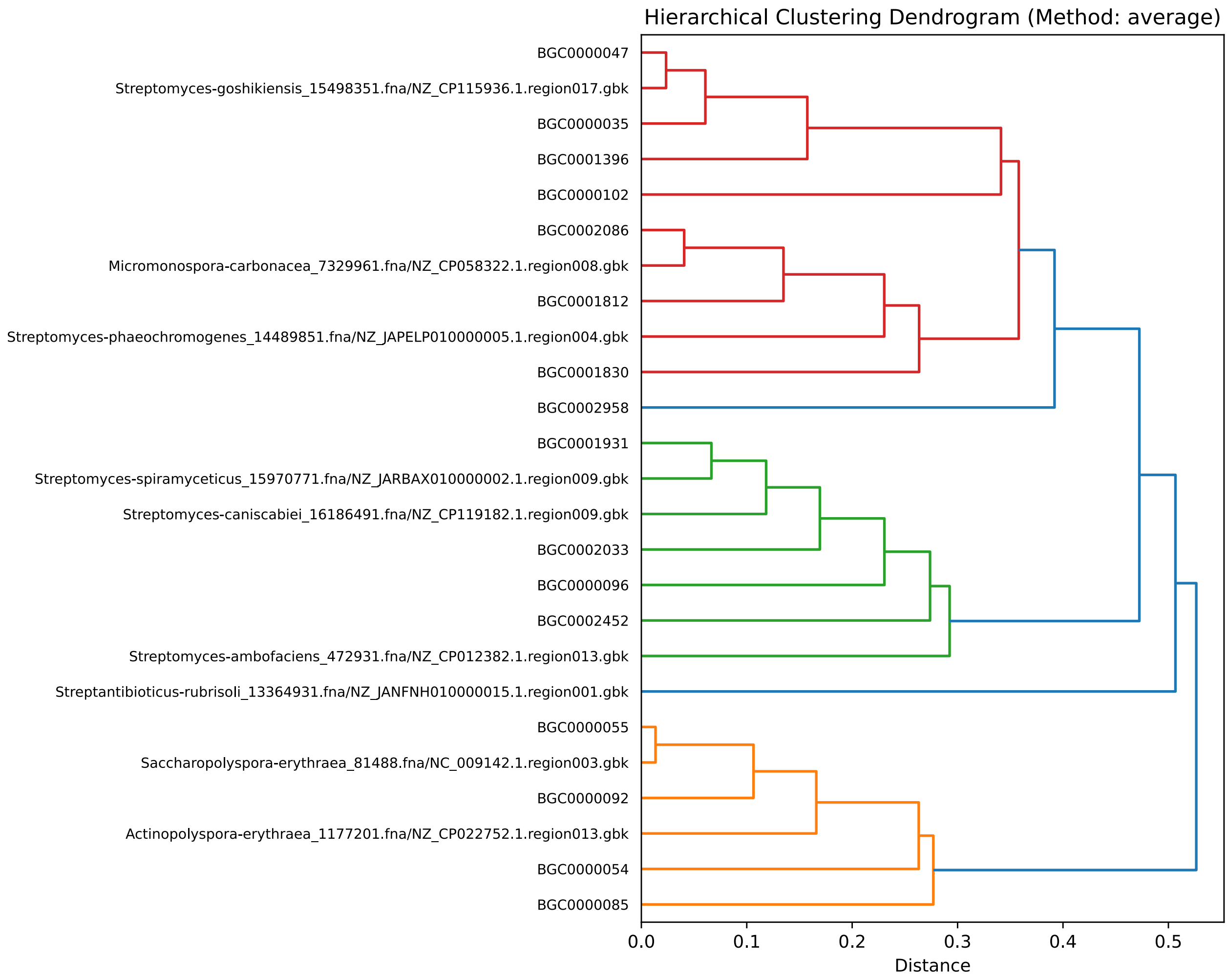

#### Glycopeptides

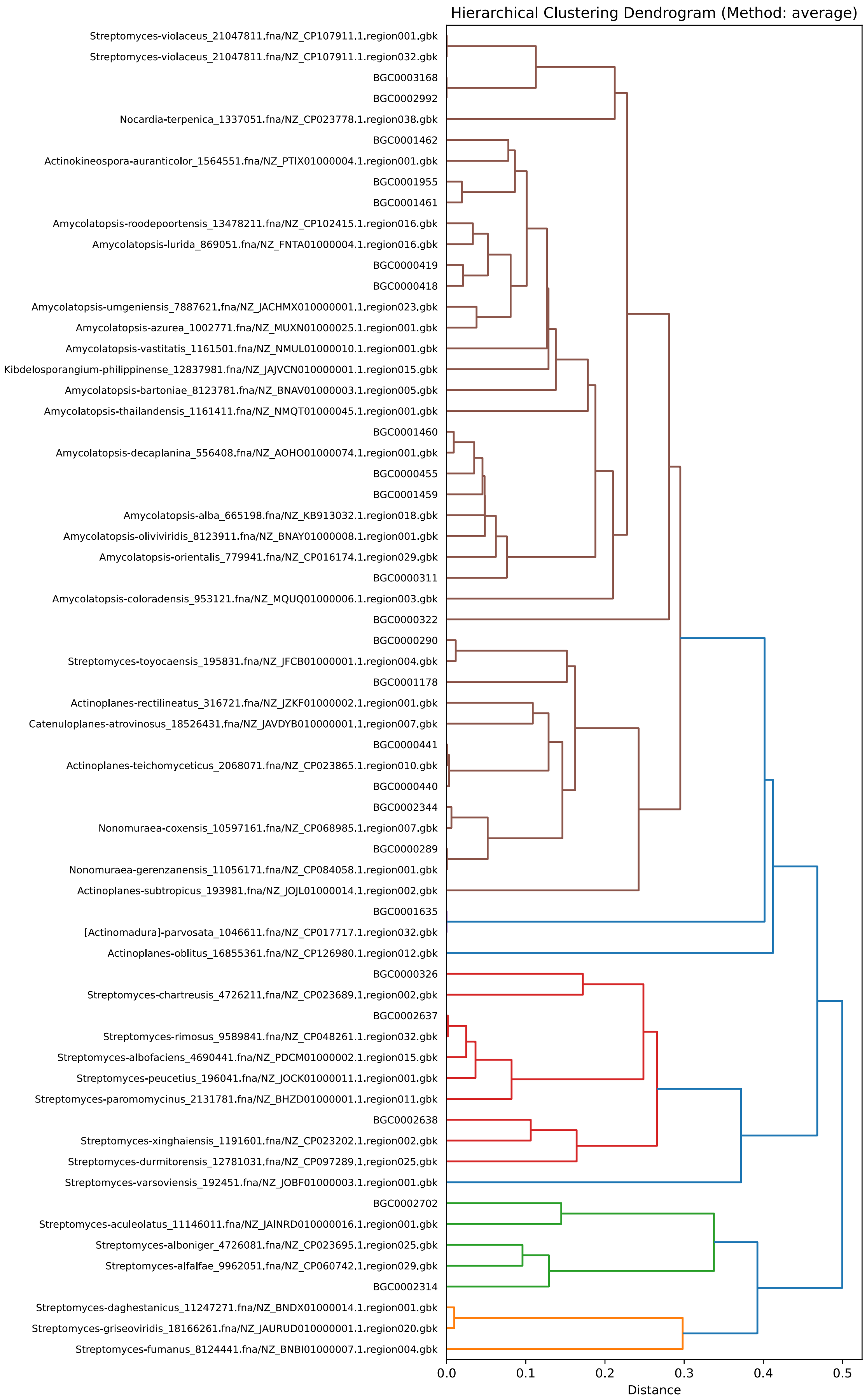

#### CDAs

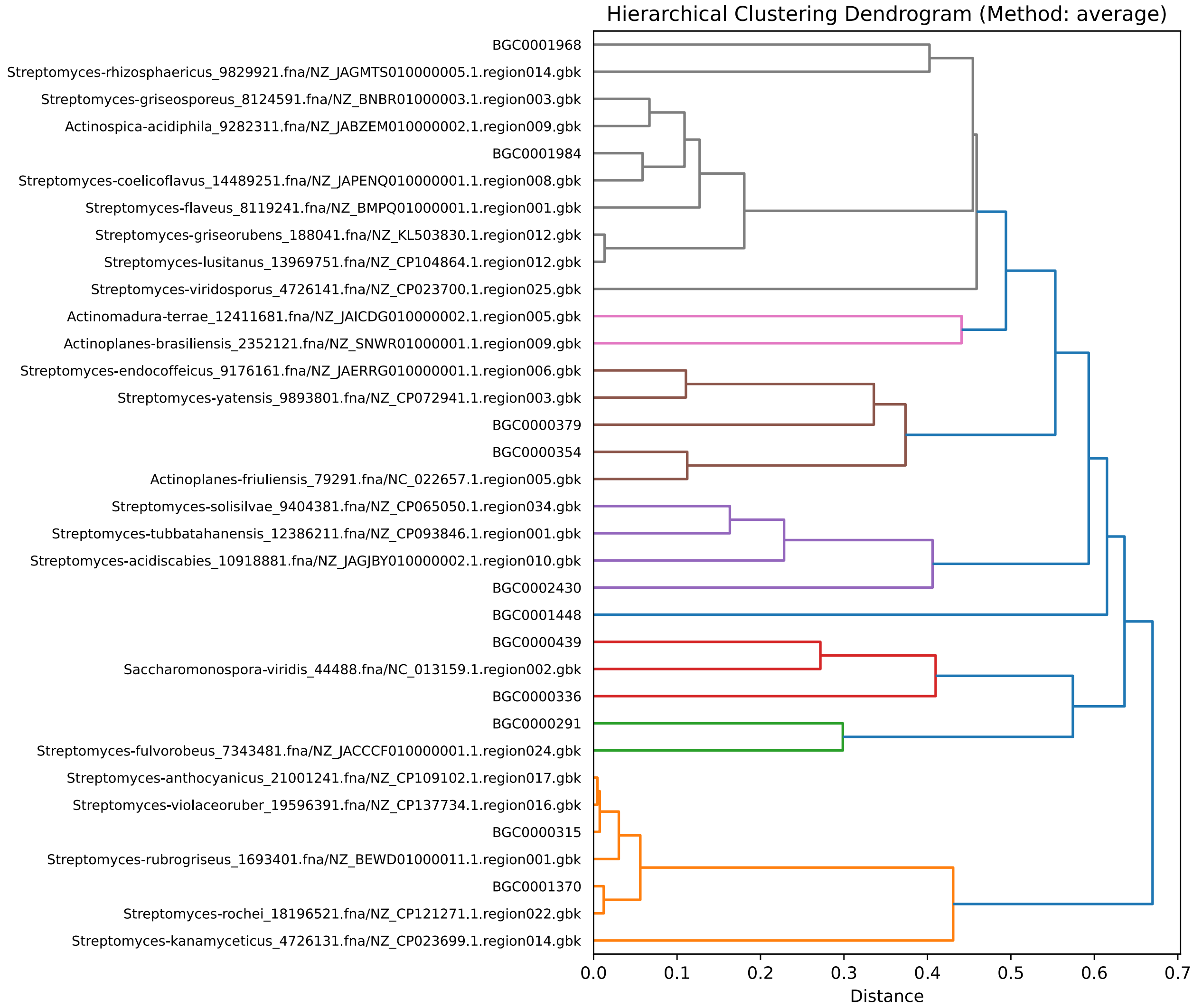
Hierarchical Clustering Dendrogram (Method: average)
BGC0001968
Streptomyces-rhizosphaericus_9829921.fna/NZ_JAGMTS010000005.1.region014.gbk
Streptomyces-griseosporeus_8124591.fna/NZ_BNBR01000003.1.region003.gbk
Actinospica-acidiphila_9282311.fna/NZ_JABZEM010000002.1.region009.gbk
BGC0001984
Streptomyces-coelicoflavus_14489251.fna/NZ_JAPENQ010000001.1.region008.gbk
Streptomyces-flaveus_8119241.fna/NZ_BMPQ01000001.1.region001.gbk
Streptomyces-griseorubens_188041.fna/NZ_KL503830.1.region012.gbk
Streptomyces-lusitanus_13969751.fna/NZ_CP104864.1.region012.gbk
Streptomyces-viridosporus_4726141.fna/NZ_CP023700.1.region025.gbk
Actinomadura-terrae_12411681.fna/NZ_JAICDG010000002.1.region005.gbk
Actinoplanes-brasiliensis_2352121.fna/NZ_SNWR01000001.1.region009.gbk
Streptomyces-endocoffeicus_9176161.fna/NZ_JAERRG010000001.1.region006.gbk
Streptomyces-yatensis_9893801.fna/NZ_CP072941.1.region003.gbk
BGC0000379
BGC0000354
Actinoplanes-friuliensis_79291.fna/NC_022657.1.region005.gbk
Streptomyces-solisilvae_9404381.fna/NZ_CP065050.1.region034.gbk
Streptomyces-tubbatahanensis_12386211.fna/NZ_CP093846.1.region001.gbk
Streptomyces-acidiscabies_10918881.fna/NZ_JAGJBY010000002.1.region010.gbk
BGC0002430
BGC0001448
BGC0000439
Saccharomonospora-viridis_44488.fna/NC_013159.1.region002.gbk
BGC0000336
BGC0000291
Streptomyces-fulvorobeus_7343481.fna/NZ_JACCCF010000001.1.region024.gbk
Streptomyces-anthocyanicus_21001241.fna/NZ_CP109102.1.region017.gbk
Streptomyces-violaceoruber_19596391.fna/NZ_CP137734.1.region016.gbk
BGC0000315
Streptomyces-rubrogriseus_1693401.fna/NZ_BEWD01000011.1.region001.gbk
BGC0001370
Streptomyces-rochei_18196521.fna/NZ_CP121271.1.region022.gbk
Streptomyces-kanamyceticus_4726131.fna/NZ_CP023699.1.region014.gbk
0.0
0.1
0.2
0.3
0.4
0.5
0.6
0.7
Distance

##### Rifamycin Like

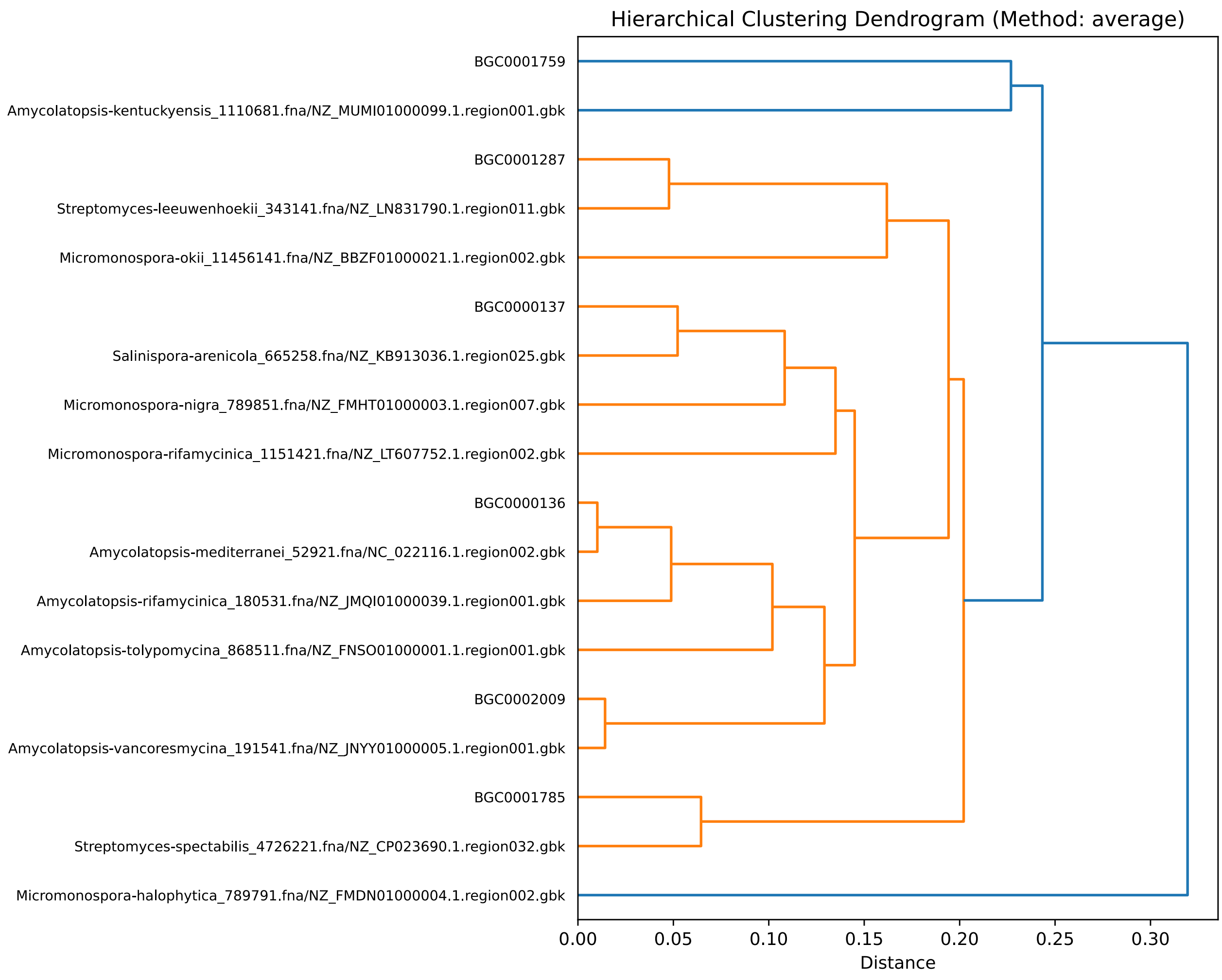
