## Supplementary material for "DiscERN: An Automated Genome Mining Tool for the Discovery of Evolutionarily Related Natural Products": BGC analysis summary

Table 1

| | Modules | Closest | Hit | Reference |
|---|---|---|---|---|
| | **CDA Filtered K4 (14/14 TP)** | | | |
| 1 | Identical | Sarpeptin | | |
| 2 | Identical | Friulimicin | | |
| 3 | Identical | CDA | | |
| 4 | V. Similar | Glycinocin | | |
| 5 | Identical | Sarpeptin | | |
| 6 | Identical | Sarpeptin | | |
| 7 | Identical | Sarpeptin | | |
| 8 | New | Cadaside | | |
| 9 | Identical | Sarpeptin | | |
| 10 | Identical | Glycinocin | | |
| 11 | Identical | CDA | | |
| 12 | Identical | CDA | | |
| 13 | Identical | CDA | | |
| 14 | Identical | Sarpeptin | | |
| | **CDA Filtered K3 (9/9 TP)** | | | |
| 1 | New | CDA | | |
| 2 | New* | Glycinocin | | |
| 3 | New | Gausemycin | | |
| 4 | New | Gausemycin | | |
| 5 | New | Ambocidin | | |
| 6 | New* | Cadaside | | |
| 7 | Identical** frag. | Gausemycin | | |
| 8 | V. Similar | Daptomycin | | |
| 9 | Identical | A54145 | | |
| | **CDA Filtered K2 (0/1 TP)** | | | |
| 1 | False Positive | N/A | | |
| | **CDA Filtered K1 (2/6 TP)** | | | |
| 1 | False Positive | Telomycin | | |
| 2 | False Positive | Omnitpeptin | | |
| 3 | False Positive | Skyllamycin | | |
| 4 | New | Ambocidin | | |
| 5 | False Positive | Cadaside | | |
| 6 | False Positive | N/A | | |
| | **CDA Unfiltered K4 (1/1 TP)** | | | |
| 1 | V. Similar | Glycinocin | | |
| | **CDA Unfiltered K3 (7/7 TP + 4 N/D partial clusters)** | | | |
| 1 | N/D, Partial | N/A | | |
| 2 | N/D, Partial | N/A | | |
| 3 | N/D, Partial | N/A | | |
| 4 | V. Similar | Glycinocin | | |
| 5 | Identical, partial | A54145 | | |
| 6 | N/D, Partial | N/A | | |
| 7 | Identical, partial | A54145 | | |
| 8 | Identical, partial | A54145 | | |
| 9 | Identical, partial | Lipopeptide 8D1 | | |
| 10 | New, partial | Friulimicin | | |
| 11 | Identical, partial | A54145 | | |
| | **CDA Unfiltered K2 (20/50 analysed, 6/13 TP + 7 N/D partial clusters)** | | | |
| 1 | Identical, partial | Friulimicin | | |
| 2 | False Positive | N/A | | |
| 3 | Identical, partial | Friulimicin | | |
| 4 | N/D, Partial | N/A | | |
| 5 | False Positive | N/A | | |
| 6 | Identical, partial | A54145 | | |
| 7 | N/D, Partial | N/A | | |
| 8 | Identical, partial | A54145 | | |
| 9 | False Positive | N/A | | |
| 10 | N/D, Partial | N/A | | |
| 11 | N/D, Partial | N/A | | |
| 12 | Identical, partial | Friulimicin | | |
| 13 | N/D, Partial | N/A | | |
| 14 | N/D, Partial | N/A | | |
| 15 | False Positive | Friulimicin | | |
| 16 | False Positive | N/A | | |
| 17 | Very Similar, partial | Glycinocin | | |
| 18 | False Positive | Glycinocin | | |
| 19 | False Positive | CDA | | |
| 20 | N/D, Partial | N/A | | |
| | **CDA Unfiltered K1 (20/214 analysed, 1/14 TP + 6 N/D partial clusters)** | | | |
| 1 | N/D, Partial | N/A | | |
| 2 | False Positive | N/A | | |
| 3 | N/D, Partial | N/A | | |
| 4 | False Positive | N/A | | |
| 5 | False Positive | N/A | | |
| 6 | False Positive | N/A | | |
| 7 | N/D, Partial | N/A | | |
| 8 | False Positive | N/A | | |
| 9 | False Positive | N/A | | |
| 10 | N/D, Partial | N/A | | |
| 11 | N/D, Partial | N/A | | |
| 12 | False Positive | N/A | | |
| 13 | False Positive | Glycinocin | | |
| 14 | Identical, partial | Taromycin | | |
| 15 | False Positive | Sarpeptin | | |
| 16 | False Positive | Cadaside | | |
| 17 | N/D, Partial | N/A | | |
| 18 | False Positive | N/A | | |
| 19 | False Positive | Friulimicin | | |
| 20 | False Positive | Glycinocin | | |
| | **Macrolides Filtered K4 (4/4 TP)** | | | |
| 1 | V. Similar | Spiramycin | | |
| 2 | New | Aldgamycin | | |
| 3 | New | Tylactone | | |
| 4 | New | Carrimycin | | |
| | **Macrolides Filtered K3 (4/5 TP)** | | | |
| 1 | Identical | Erythromycin | | |
| 2 | New | Carrimycin | | |
| 3 | False Positive | Chalcomycin | | |
| 4 | Identical | Erythromycin | | |
| 5 | Identical | Spiramycin | | |
| | **Macrolides Filtered K2 (3/3 TP)** | | | |
| 1 | New | Rosamicin | | |
| 2 | Identical | Erythromycin | | |
| 3 | Identical | Lankamycin | | |
| | **Macrolides Filtered K1 (0/2 TP)** | | | |
| 1 | False Positive | Aldgamycin | | |
| 2 | False Positive | Aldgamycin | | |
| | **Macrolides Unfiltered K3 (2/2 TP)** | | | |
| | Identical, partial | Spiramycin | | |
| | Identical | Lankamycin | | |
| | **Macrolides Unfiltered K2 ( 3/4 TP + 5 N/D partial** | | | |
| 1 | New | Chalcomycin | | |
| 2 | New | Rosamicin | | |
| 3 | New | Rosamicin | | |
| 4 | N/D, Partial | Spiramycin | | |
| 5 | N/D, Partial | Rosamicin | | |
| 6 | N/D, Partial | Mycinamicin | | |
| 7 | False Positive | Carrimycin | | |
| 8 | N/D, Partial | Leucomycin | | |
| 9 | N/D, Partial | Chalcomycin | | |
| | **Macrolides Unfiltered K1 (0/2 TP + 5 N/D Partial)** | | | |
| 1 | N/D, Partial | Rosamicin | | |
| 2 | N/D, Partial | Lankamycin | | |
| 3 | N/D, Partial | Ibomycin | | |
| 4 | N/D, Partial | Carrimycin | | |
| 5 | False Positive | Polyoxypeptin | | |
| 6 | False Positive | Efomycin | | |
| 7 | N/D | Dehydro-chalcomycin | | |
| | **Rifamycin Filtered K3 (1/1 TP)** | | | |
| 1 | Identical | Chaxamycin | | |
| | **Rifamycin Filtered K1 (1/1 TP)** | | | |
| 1 | Identical | Kanglemycin | | |
| | **Rifamycin Unfiltered K4 (9/9 TP)** | | | |
| 1 | Identical, partial | Rifamycin | | |
| 2 | Identical | Kanglemycin | | |
| 3 | Identical | Rifamycin | | |
| 4 | Identical | Kanglemycin | | |
| 5 | Identical* | Kanglemycin | | |
| 6 | V. Similar | Rifamycin | | |
| 7 | Identical | Rifamycin | | |
| 8 | New | Rifamycin | | |
| 9 | Identical | Rifamycin | | |
| | **Rifamycin Unfiltered K3 (2/2 TP)** | | | |
| 1 | Identical | Streptovaricin | | |
| 2 | Identical, partial | Rifamycin | | |
| | **Rifamycin Unfiltered K2 (3/3 TP)** | | | |
| 1 | Identical, partial | Rifamycin | | |
| 2 | N/D, Partial | Streptovaricin | | |
| 3 | Identical, partial | Rifamycin | | |
| 4 | Identical, partial | Chaxamycin | | |
| | **Rifamycin Unfiltered K1 (3/11 TP + 13 N/D Partial)** | | | |
| 1 | False Positive | N/A | | |
| 2 | False Positive | Hygrocin | | |
| 3 | Identical, partial | Streptovaricin | | |
| 4 | N/D, Partial | N/A | | |
| 5 | False Positive | Rubradirin | | |
| 6 | N/D, Partial | Chaxamycin | | |
| 7 | False Positive | Rubradirin | | |
| 8 | False Positive | Microansamycin | | |
| 9 | False Positive | N/A | | |
| 10 | N/D, Partial | Napthomycin | | |
| 11 | N/D, Partial | N/A | | |
| 12 | N/D, Partial | Rifamycin | | |
| 13 | N/D, Partial | N/A | | |
| 14 | N/D, Partial | N/A | | |
| 15 | N/D, Partial | N/A | | |
| 16 | False Positive | rifamycin | | |
| 17 | N/D, Partial | N/A | | |
| 18 | False Positive | Kanglemycin | | |
| 19 | N/D, Partial | N/A | | |
| 20 | Identical, partial | Rifamycin | | |
| 21 | N/D, Partial | N/A | | |
| 22 | Identical, partial | Rifamycin | | |
| 23 | N/D, Partial | N/A | | |
| 24 | N/D, Partial | N/A | | |
| | **Glycopeptides Filtered K4 (20/36 analysed, 20/20 TP)** | | | |
| 1 | Identical | Rimomycin | | |
| 2 | New | Ristomycin | | |
| 3 | Identical | Ristomycin | | |
| 4 | Identical | UK-68,597 | | |
| 5 | Identical | Ristomycin | | |
| 6 | Identical | Keratinimicin | | |
| 7 | Identical | Decaplanin | | |
| 8 | Identical | Rimomycin | | |
| 9 | Identical | Teicoplanin | | |
| 10 | Identical | misaugamycin | | |
| 11 | Identical | Rimomycin | | |
| 12 | Identical | Teicoplanin | | |
| 13 | Identical | A40926 | | |
| 14 | Identical | Kistamycin | | |
| 15 | Identical | Ristomycin | | |
| 16 | Identical | Ristomycin | | |
| 17 | Identical | Rimomycin | | |
| 18 | Identical | A50926 | | |
| 19 | Identical | Avoparcin | | |
| 20 | Identical | Decaplanin | | |
| | **Glycopeptides filtered K3 (6/6 TP)** | | | |
| 1 | New | Cobromycin | | |
| 2 | New | Cobromycin | | |
| 3 | Identical | Cobromycin | | |
| 4 | New | Cobromycin | | |
| 5 | Identical | Cobromycin | | |
| 6 | Identical | UK-68,597 | | |
| | **Glycopeptides filtered K2 (1/1 TP)** | | | |
| | New | Cobromycin | | |
| | **Glycopeptides unfiltered K4 (3/3 TP)** | | | |
| 1 | Identical, partial | misaugamycin | | |
| 2 | Identical | rimomycin | | |
| 3 | Identical, partial | Kistamycin | | |
| 4 | New, partial, fragmented | Cobromycin | | |
| | **Glycopeptides unfiltered K3 (7/7 TP + 5 N/D Partial)** | | | |
| 1 | New, partial | Cobromycin | | |
| 2 | Identical, partial | Kistamycin | | |
| 3 | Identical, partial | rimomycin | | |
| 4 | N/D, Partial | N/A | | |
| 5 | N/D, Partial | N/A | | |
| 6 | New | Cobromycin | | |
| 7 | N/D, Partial | N/A | | |
| 8 | Identical, partial | Cobromycin | | |
| 9 | N/D, Partial | N/A | | |
| 10 | New | Cobromycin | | |
| 11 | New | Cobromycin | | |
| 12 | N/D, Partial | N/A | | |
| | **Glycopeptides unfiltered K2 (20/53 analysed, 2/5 TP + 15 N/D Partial)** | | | |
| 1 | N/D, Partial | N/A | | |
| 2 | N/D, Partial | N/A | | |
| 3 | Identical, partial | Cobromycin | | |
| | N/D, Partial | N/A | | |
| 4 | False Positive | Cobromycin | | |
| 5 | N/D, Partial | N/A | | |
| 6 | N/D, Partial | N/A | | |
| 7 | N/D, Partial | N/A | | |
| 8 | N/D, Partial | N/A | | |
| 9 | N/D, Partial | N/A | | |
| 10 | False Positive | N/A | | |
| 11 | N/D, Partial | N/A | | |
| 12 | N/D, Partial | N/A | | |
| 13 | N/D, Partial | N/A | | |
| 14 | N/D, Partial | N/A | | |
| 15 | New, partial | Cobromycin | | |
| 16 | N/D, Partial | N/A | | |
| 17 | N/D, Partial | N/A | | |
| 18 | N/D, Partial | N/A | | |
| 19 | False Positive | feglymycin | | |
| 20 | N/D, Partial | N/A | | |
| | **Glycopeptides unfiltered K1 (20/198 analysed, 0/16 TP + 4 N/D Partial)** | | | |
| 1 | False Positive | N/A | | |
| 2 | N/D, Partial | N/A | | |
| 3 | False Positive | N/A | | |
| 4 | False Positive | N/A | | |
| 5 | False Positive | N/A | | |
| 6 | N/D, Partial | N/A | | |
| 7 | N/D, Partial | N/A | | |
| 8 | False Positive | N/A | | |
| 9 | False Positive | N/A | | |
| 10 | False Positive | N/A | | |
| 11 | False Positive | N/A | | |
| 12 | False Positive | | | |
| 13 | N/D, Partial | N/A | | |
| 14 | False Positive | coprisamide | | |
| 15 | False Positive | N/A | | |
| 16 | False Positive | N/A | | |
| 17 | False Positive | N/A | | |
| 18 | False Positive | feglymycin | | |
| 19 | False Positive | feglymycin | | |
| 20 | False Positive | Kistamycin | | |
